# Interleukin-3 as a Potential Bone Anabolic Agent in treating Postmenopausal Osteoporosis

**DOI:** 10.1101/2025.07.12.664485

**Authors:** Vikrant Piprode, Shubhanath Behera, Juilee Karhade, Garima Pandey, Arpita Prasad, Krishna Ezhuthachan, Mohan R. Wani

## Abstract

Postmenopausal osteoporosis (PMO), a silent disorder caused due to estrogen deficiency, is characterized by loss of bone mass and low bone mineral density. Despite advancements in treatment, current medications often fail to fully restore bone integrity and prevent fragility fractures in the osteoporotic individuals. Thus, there is a high demand of novel therapeutic modalities in dealing with osteoporosis. Interleukin-3 (IL-3), a cytokine secreted by activated T cells, emerges as a promising therapeutic candidate. The dual role of IL-3 in inhibiting osteoclast differentiation and promoting osteoblast differentiation, offers protection against bone and joint degeneration in arthritic mice. However, its role in osteoporosis is not yet delineated. Therefore, our investigation focuses on elucidating the role of IL-3 in PMO, employing both prophylactic and therapeutic strategies in ovariectomized mice, mimicking human PMO. In our study, at the onset of osteoporosis, IL-3 treatment effectively preserved trabecular bone architecture and enhanced bone mineral density in OVX mice, particularly notable in therapeutic interventions where fully developed osteoporosis was addressed. Notably, in preventive measures, IL-3 inhibited osteoclast differentiation, thereby suppressing bone resorption, while in therapeutic approaches, it enhanced osteoblast differentiation, promoting bone formation. Irrespective of gender specific, microCT analysis of IL-3^-/-^ (KO) mice showed reduced trabecular bone development compared to its respective wild type (WT) mice, highlighting essential role of IL-3 in skeletal integrity. Moreover, IL-3 increased Treg cells population while inhibiting B cell lymphopoiesis in ovariectomized mice with no adverse effects on hematopoiesis or vital organs. In conclusion, our findings collectively underscore the potential of IL-3 as a novel therapy for PMO, offering insights into its mechanisms of action and clinical applications.

**One Sentence Summary:** IL-3 has a dual action on bone: anti-osteoclastic and pro-osteoblastic in OVX mice, thus can be a novel therapeutic drug in treating postmenopausal osteoporosis.

## Introduction

Postmenopausal osteoporosis (PMO), a silent condition that occurs after menopause, is primarily attributed to lack of estrogen, which results in substantial bone loss (*1*). It is characterized by reduced bone mass and density, leading to heightened bone fragility and an increased risk of fractures (*2*). It affects nearly 200 million people globally and is often called a silent disease since it often goes unnoticed until a bone fracture occurs (*3*). Although menopause is natural, but not all postmenopausal women suffer from osteoporosis as those who attain peak bone mass during their growth years are less likely to develop osteoporosis, even after menopause. In general, insufficient peak bone mass, excessive bone resorption, or impaired bone formation post-menopause, all contribute to osteoporotic bone loss (*4*).

Current FDA approved drugs to treat PMO include anti-resorptives drugs, bone-forming synthetic analogs of PTH (Teriparatide and Abaloparatide), and antibodies like Denosumab and Romosozumab, which are antibodies to inhibits osteoclast formation and enhance bone formation respectively. However, each of these drugs is either anti-osteoclast or pro-osteoblast, and all anabolic agents are sold with a black box warning to indicate deleterious side effects. An ideal osteoporosis treatment drug should exhibit both anti-osteoclastic and pro-osteoblastic characteristics, demonstrate anti-fracture efficacy across diverse skeletal sites (spine, non-vertebral areas, and hip), offer convenient administration and treatment intervals, be cost-effective, and have minimal to no side effects.

We have documented that a cytokine, interleukin-3 (IL-3), primarily secreted by activated T cells, is a potent inhibitor of osteoclast differentiation and bone resorption in vitro in rodents(*5–8*).Similarly, Gupta et al., 2010Gupta et al., 2010confirmed inhibitory effect of IL-3 on RANKL-induced osteoclast differentiation in osteoclast precursors from the blood monocytes and bone marrow cells of individuals with osteoporosis. Further research revealed that IL-3 enhances osteoblast differentiation and bone mineralization in both in vitro and in vivo (*10*), implying its potential to inhibit abnormal bone resorption and increase osteoblast numbers responsible for bone formation. Also, IL-3 primed osteoblasts are protected from the negative actions of TNF-α (*11*). Additionally, IL-3 prevents the onset of inflammatory arthritis as well as osteoarthritis in mice and protects joint cartilage and bone integrity (*6, 12*). Yet, the role of IL-3 in pathological disease conditions of PMO remains unclear. To address this issue, we established an in vivo mouse model for osteoporosis and examined the potential preventive and therapeutic effects of IL-3 treatment on OVX-induced osteoporotic mice. To further corroborate our findings, we also conducted an investigation into the impact of IL-3 deficiency on bone architecture, assessing the bone structure in IL-3 KO mice.

To evaluate the efficacy and safety profile of any treatment, it is essential to study its effects across different stages of disease progression, as pathology can vary significantly from early to late stages. In PMO, where bone loss becomes more severe over time, it is critical to investigate both prophylactic (before the initiation of bone loss) and therapeutic (after bone loss is evident) interventions. Prophylactic studies allow for assessing whether IL-3 can inhibit ovariectomy-induced bone loss by targeting bone-resorbing osteoclasts. This approach focuses on the anti-osteoclastic potential of IL-3, aiming primarily to prevent bone resorption rather than stimulate bone formation. In contrast, therapeutic studies evaluate whether IL-3 can restore bone architecture that has already been compromised by targeting bone-forming osteoblasts. In this context, the anabolic effects of IL-3 are examined to determine if it can enhance bone formation. By distinguishing between these two stages, the overall efficacy of IL-3 in both preventing and reversing bone loss can be comprehensively assessed.

## Material and methods

### Animals

Adult female C57BL/6J mice (8-10 weeks) were used for the development of animal model of osteoporosis. IL-3 knock out (KO) mice (B6.129S2(B10)-Il3tm1Tyb/J) were purchased from Jackson laboratory. Both male and female IL-3 KO mice of different age group were used for hematological study and characterization of skeletal micro-architecture. All mice were sourced from the Experimental Animal Facility of the National Centre for Cell Science (NCCS), Pune, India. All the animals had access to food and water ad libitum and were euthanized via CO2 asphyxiation. All protocols involving animal use were approved by an Institutional Animal Ethics Committee (IAEC), NCCS.

### Development of mouse model of osteoporosis by ovariectomy and IL-3 administration strategy

Animal model of osteoporosis was developed according the previous reports (*13*). To develop osteoporotic mice model eight weeks old female C57BL/6J mice were used. The mice were divided into three groups with 10-12 mice in each-SHAM (as control), OVX (ovariectomized), and OVX+IL-3 (OVX mice injected with IL-3). In brief, the mice were anaesthetized using a combination of xylazine (0.01 mg/g body weight) and ketamine (0.1 mg/g body weight). To create access into the abdomen area, the hair was shaved using a razor on one side of the flank (between ribcage and hip) and swabbed with Betadine solution. An incision (approx. 1 cm) on the skin was first made to expose the underling muscles. A smaller incision (<1 cm) in the muscle layer was made to allow entry into the peritoneal cavity. The ovaries were identified in the fat pad adjacent to the perinephric region and were completely excised. The muscle incision was closed using catgut sutures (absorbable) followed by skin incision with non-absorbable sutures. The same procedure was followed for sham operations except that the ovaries were identified but not removed. To study the effect of IL-3 in osteoporosis two different strategies were followed: one was prophylactic study and another one was therapeutic study. In prophylactic study, IL-3 (1 μg/day) was injected i.p. in OVX mice (OVX+IL-3 mice group), from day 8 to day 37 of post-ovariectomy. At day 38 all the mice were euthanized; long bones were excised. Both femur and tibia were subjected to micro-computed tomography (μ-CT) analysis to study the bone phenotype. In therapeutic study IL-3 (1 μg/day) was injected i.p. in OVX (OVX+IL-3 mice group), mice after the development of osteoporotic model i.e., from day 30 to day 59 of post-ovariectomy. At day 60 all the mice were euthanized; long bones were excised. Both femur and tibia were subjected to μ-CT analysis to study the bone phenotype.

### Isolation of osteoclast precursors (OCPs) from mice bone marrow and in vitro osteoclast differentiation

Bone marrow cells were isolated from C57BL/6J mice of 8-10 weeks as described earlier (*7*). Mice were euthanized and long bones were separated, the epiphyseal end of each bone was cut and the marrow was flushed. The cells were dispersed, washed and resuspended in the alpha-modified MEM medium (Sigma Aldrich; catalogue #M8042) were incubated with recombinant mice M-CSF (Biolegend; catalogue # 576404,) (15 ng/ml) in a 25-cm^2^ flask from one mouse. After 24 hrs, the non-adherent cells were removed, washed and subjected to RBC lysis. The mononuclear OCPs at a density of 5 X 10^4^ cells/well were plated in a 96-well plate. Then the cells were treated with recombinant mouse M-CSF (30 ng/ml), recombinant mouse RANKL (Peprotech; catalogue #315-11-10) (40 ng/ml) without or with recombinant mouse IL-3 (Peprotech; catalogue # 213-13-10) (10 ng/ml). Cultures were half fed on every 3^rd^ day by replacing half of culture media with fresh media and factors. On day 4 culture supernatant was collected from each well and cells were fixed with 10% formalin. The fixed cells and the supernatant were subjected to TRAP staining.

### Characterization of osteoclasts by TRAP staining

To assess tartrate-resistant acid phosphatase (TRAP) qualitative assay cells were washed with 1X PBS and formalin fixed for 10 minutes. A freshly prepared TRAP staining mixture was used in this procedure. Pararosaniline chloride (40 mg/ml) was mixed with a 25% concentrated HCl solution and heated in a glass beaker at 60°C for 10 minutes until dissolved. A stock buffer solution was made by dissolving sodium acetate (11.7 mg/ml), sodium barbitone (29.4 mg/ml), and tartaric acid (10 mg/ml) in distilled water. The substrate, naphthol AS-BI phosphate (25 mg), was dissolved in 2.5 ml of N, N dimethylformamide in a glass tube. Sodium nitrite solution (40 mg/ml) was freshly prepared in distilled water right before use. The pH was maintained to 4.7 using sodium hydroxide solution. and the mixture was then filtered. The fixed cells were stained with above mixture and incubated for 20 minutes at room temperature and subsequently counterstained with Mayer’s hematoxylin. Finally, the stained cells were examined under a bright field microscope, and TRAP-positive cells, which appeared pink, were counted, with a focus on cells with three or more nuclei.

For intracellular quantitative TRAP assay, the osteoclast cultures were first rinsed with 1X PBS, and then were fixed with 10% formalin for 10 minutes. After fixation, they were washed with absolute ethanol for 1 minute. Next, the fixed cells were incubated with a solution containing 50 mM citrate buffer (pH 4.5), 10 mM sodium tartarate, and 5 mM 4-nitrophenyl phosphate (PNPP) for 60 minutes at 37°C. The enzymatic reaction was stopped by adding 1N NaOH, and the absorbance was measured at 415 nm using a microplate reader. For quantitative extracellular TRAP assay, the culture supernatant was subjected to the same protocol as described above using pNPP substrate.

### Micro-computed tomography (μ-CT)

To study the bone histomorphometry of femur and tibia μ-CT was performed using Skyscan 1276 μ-CT Scanner (Sky Scan, Ltd., Kartuizersweg, Konitch, Belgium). Left femora and tibia were excised from the animals after sacrifice, cleaned of soft tissue, and fixed in 10% formalin for 48 h. Bone sample scanning was performed at 50 kV, 100 μA using a 0.5 mm aluminium filter. All the projections were collected at a resolution of 7 μm/pixel and reconstructed using NRecon software. CTAnalyser software was used for 3D morphometric analysis of trabecular and cortical bone indices which are as follows: bone volume (BV), bone volume/tissue volume (BV/TV), trabecular thickness (Tb.Th), trabecular number (Tb.N), connectivity density (Conn.D), trabecular space (Tb.Sp), cortical thickness (Ct.Th), cortical bone volume/tissue volume (Ct.BV/TV) and Ct.medullary volume. Bone mineral density (BMD) of trabecular as well as cortical bone was also analysed separately. 3D models of trabecular and cortical bone were developed using CTvol software.

### Blood collection, CBC and serum biochemical analysis

Blood samples from each mouse were collected via retro-orbital sinus bleeding using a micro-hematocrit capillary. Blood samples were either collected in Vacutainer SSTTM II advance (for serum collection) or in EDTA (1.8 mg/ml) containing 1.5 ml Eppendorf tube (for complete blood count-CBC test) or lithium-heparin containing tube. For serum isolation blood sample were kept at room temperature for 30 minutes and then centrifuged at 2500 rpm for 10 minutes. Using a sterile pipette, the serum was carefully collected from the top layer of the separated blood (the liquid portion) and transfer it to sterile microcentrifuge tubes. All the serum samples were stored at -80°C till further use. For CBC test EDTA containing of blood samples (100 μl) were run through Celltac α MEK-6550K automated hematology analyzer using mouse species program. For general health profiling of IL-3 KO mice, 500 μl of blood was collected in lithium heparin tubes provided by manufacturer. The tubes were gently mixed by inverting for 30 seconds. The blood samples were then centrifuged at 12,000 rcf for 120 seconds to separate plasma. The plasma samples were collected and analysed using VetTest® chemistry analyzer following the provided instruction manual.

### Flow cytometry

To study the distribution of immune cells, peritoneal fluid, spleen, lymph nodes and bone marrow were dissected from all the mice and kept in complete RPMI-1640 media separately. Briefly single cell suspensions were prepared by mashing the spleen and lymph-nodes separately through the cell strainer into the petri dish and bone marrow was flushed from femur and tibia. Subsequently, the cell suspensions were centrifuged at 1500 rpm for 5 minutes, after which 1 ml of RBC lysis buffer (Invitrogen™ eBioscience™; catalogue #00-4333-57) was added. Cells were washed thoroughly with complete RPMI-1640 media (Gibco™; catalogue #11875093) and re-suspended with 1 ml complete RPMI-1640 media. Viable cell count was determined using trypan blue and haemocytometer. For flow cytometry experiment 106 cells were surface-stained with fluorochrome-conjugated antibodies and incubated for 30 minutes on ice. After thorough washing with complete RPMI, cells were fixed-permeabilized with Foxp3 permeabilization buffer (Invitrogen; eBioscience™ Foxp3/Transcription Factor Staining Buffer Set catalogue #00-5523-00) for 30 minutes on ice. Intracellular staining was carried out using fluorochrome-conjugated antibodies in permeabilization buffer for 45 minutes on ice. The antibodies used are: PE-Cyanine7 labelled anti-mouse T-bet mAb (clone 4B10) (Invitrogen; catalogue # 25-5825-82), Alexa Fluor 488 labelled anti-mouse Gata-3 mAb (clone TWAJ) (Invitrogen; catalogue # 53-9966-42), APC labelled anti-mouse Foxp3 mAb (clone FJK-16s) (Invitrogen; catalogue #17-5773-82), PE labelled anti-mouse Rorγt mAb (clone AFKJS-9) (Invitrogen; catalogue #12-6988-82), from eBioscience. Fluorochrome Percp.cy5.5 labelled anti-mouse CD4 mAb (clone RM4.5) (BD biosciences; catalogue #550954), Percp.cy5.5 labelled anti-mouse CD19 (clone 1D3) (BD biosciences; catalogue #561113), FITC labelled anti-mouse CD45R/B220 (clone RA3-6B2) (BD biosciences; catalogue # 553087)and PE labelled anti-mouse IgM mAb (clone R6-60.2) (BD biosciences; catalogue # 553409) were purchased from BD bioscience. Post-staining, cells were washed with permeabilization buffer and analyzed using BD FACSCanto^TM^ II cytometer.

### ELISA

The serum level of CTX-I (CUSABIO; CSB-E12782m) and OCN (CUSABIO ; CSB-E06917m) was assessed by ELISA according to manufacturers’ instructions.

### RNA isolation from bone tissue and qPCR-based gene expression

Total RNA was isolated from bone tissue without marrow by grinding it in liquid nitrogen in a pestle and mortar and RNA isolation was performed by TRizol method developed by Chomczynski and Sacchi in 1987. Subsequently, RNA was quantitated suing NanoDrop (NanoDrop Technologies, USA) and 1 μg of total RNA was used to prepare cDNA using M-MLV Reverse Transcriptase (Invitrogen; catalogue # 28025013). Expression of osteoclast and osteoblasts specific markers gene expression was assesses using qPCR. Each reaction consisted of 2X Power Up^TM^ SYBR green master mix (Invitrogen; catalogue #A25742 and forward and reverse primers of 100 pM each. The universal program of 40 cycles in the StepOne Plus system (Applied Biosystems) was used. Each cycle involved denaturation at 94°C for 15 seconds, followed by primer annealing and extension at 60°C for 60 seconds. Finally, a melt curve analysis was performed. The data obtained were analyzed using the comparative 2^-ΔΔCt^ method. List of primers used are summarized (**Table S1**).

### Genotyping

To confirm the IL-3 gene knock out, genotyping was conducted by PCR of tail lysate with provided primers. In brief mice tail sample was collected and kept at 55°C for overnight digestion with lysis buffer. On next day genomic DNA was extracted using phenol chloroform isoamyl alcohol method and quantified using NanoDrop. To perform PCR, the following steps were followed: In PCR tube 10 μl of EmeraldAmp GT PCR master mix, 0.5 μl mutant forward, 0.5 μl common primer, 0.5 μl wild type reverse primer and 7.5 μl of nuclease free water were added along with 1 μl DNA template and mixed. The PCR reaction was then carried out using an Eppendorf thermal cycler. The PCR reaction involved 30 cycles of denaturation at 98°C for 10 seconds, primer annealing at 50°C for 30 seconds and extension at 72°C for 1 minutes.

The primers are as follows: IL-3 Mutant Forward -AAGGGGCCACCAAAGAAC; IL-3 Common-GGGTTTTTGGCATCTTGGT; IL-3 Wild type Reverse-CCAGGGAGATGAGATCCAGA.

### Histology

The hind limbs of mice, including the intact femur and tibia, were isolated and incubated in 10% formalin for 48 hours for fixation. Subsequently, the bones underwent a decalcification process using 12.5% EDTA solution at pH 7.0, which lasted for a period of three weeks. After decalcification, the bones were embedded in paraffin wax. Longitudinal sections with a thickness of 8 μm were then obtained from the embedded bones using a microtome. Vital organs, including liver, heart and kidneys were also harvested, formalin-fixed and paraffin-embedded. Finally, the sections were stained with hematoxylin and eosin (H&E) to facilitate microscopic examination and analysis. A pathologist from a local hospital examined the tissues for signs of any pathology.

### Statistical analysis

The results are presented as the mean ± standard error of the mean (SEM) for different experimental groups. Statistical variances between groups were assessed using one way ANOVA followed by Boneferroni’s correction for multiple comparisons.

## Results

### IL-3 inhibits osteoclastogenic potential of bone marrow-derived OC precursors from OVX mice in vitro

Our group was the first to report the anti-osteoclastogenic action of IL-3 in vitro on OC precursors derived from healthy mice, rats, and human (*5–9, 14*). However, all these studies demonstrated the anti-osteoclastic potential in vitro from physiological healthy OC precursors, which did not experience any abnormal signals from surrounding cells nor were isolated from any known pathological conditions of bones. Also, compelling evidences suggest OC precursors from PMO individuals have higher osteoclast differentiation potential and function, suggesting presence of we have coined the term “pathogenic OC precursors”. Thus, we first determined the anti-osteoclastic potential of IL-3 on pathogenic OC precursors derived from osteoporotic mice. This was achieved by ovariectomizing mice and isolating bone marrow-derived OC precursors after significant bone loss has occurred, and driving RANKL-induced OC differentiation **(Fig. 1A)**. Notably, IL-3 caused complete inhibition of multinuclear TRAP^+^ cells in cultures from both SHAM and OVX mice **(Fig. 1B).** Quantitatively, IL-3 significantly reduced both the intracellular (**Fig. 1C**) and extracellular TRAP activity (**Fig. 1D**), as assessed by pNPP colorimetric assay. These findings indicate a potent anti-osteoclastic action of IL-3 on pathogenic OC precursors derived from OVX mice.

**Figure 1.**
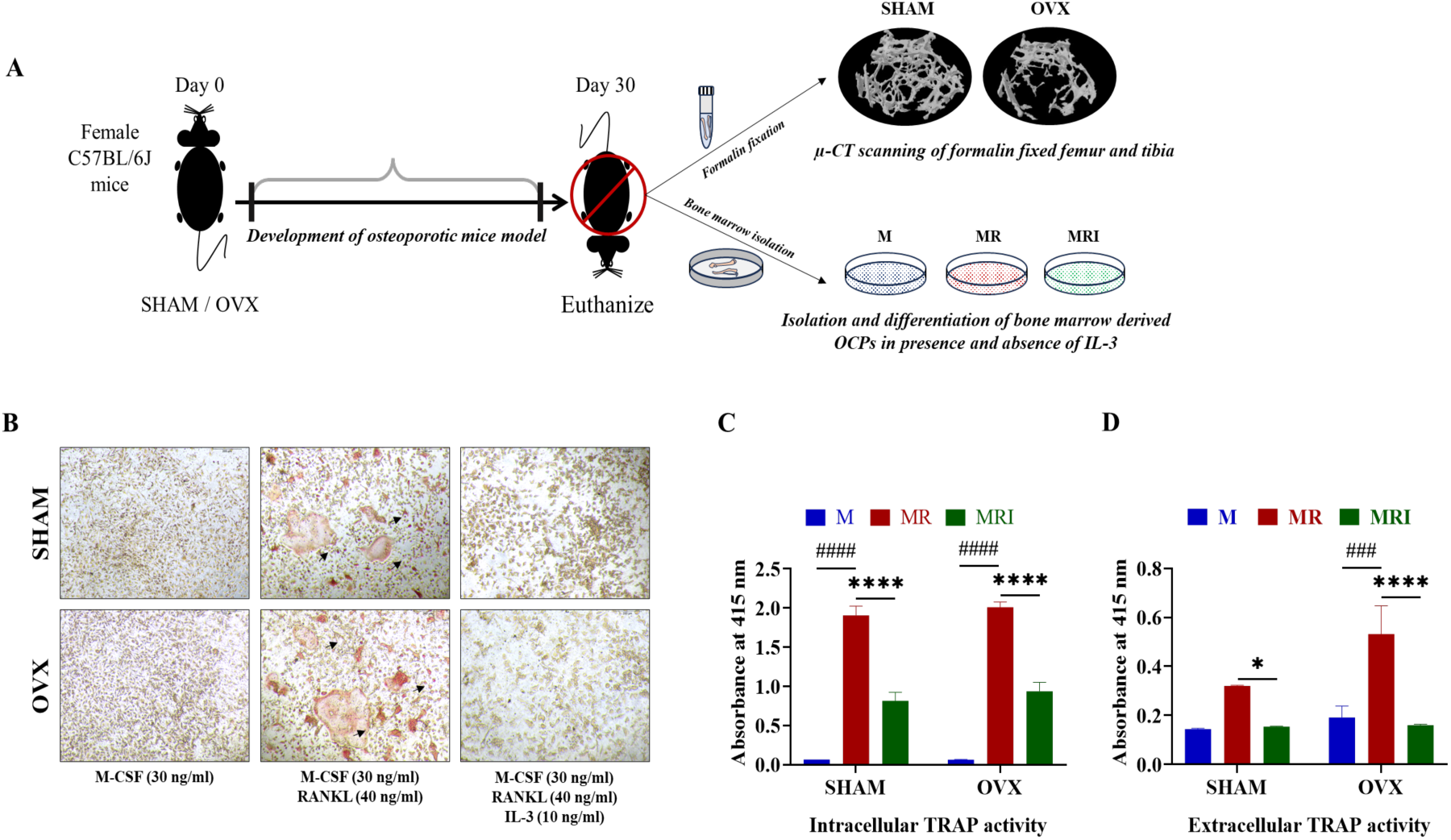
IL-3 inhibits RANKL-induced differentiation of pathogenic OC precursors derived from OVX mice in vitro. **(A)** 8-10 weeks old mice were ovariectomized and OC precursors were isolated, 30 days post-ovariectomy, from bone marrow of SHAM and OVX mice and cultured in vitro with M-CSF (M, 30 ng/ml) with or without RANKL (R, 40 ng/ml) and in absence or presence of IL-3 (10 ng/ml). After 72 hours, cells were (**B**) TRAP stained to assess the multinuclear osteoclast formation. Black arrows highlighted the TRAP^+^ mature osteoclasts (Magnification 10X; scale 200µm). In a similar experiment **(C)** intracellular and **(D)** extracellular TRAP activity was measured colorimetrically. Data presented is a representation of three independent experiments. *, p ≤ 0.05; ****, p ≤ 0.0001 vs. MR group.

### Prophylactic action of IL-3 prevents bone loss in OVX mice

As we have proven consistently in different species that IL-3 is a strong anti-osteoclastic agent, and thus we believe it could be used as a potential anti-osteoclastic agent in preventing bone loss in PMO. Therefore, we evaluate the prophylactic effects of IL-3 on bone loss in response to estrogen deficiency. We employed ovariectomy-induced bone loss mouse model using 8-10-week-old C57BL/6J female mice. This is the most widely used murine model typically mimicking the bone loss upon estrogen deficiency in postmenopausal women. In this strategy-I of IL-3 treatment, on day 8 post-OVX surgery, mice were i.p. injected with a single dose of IL-3 (1 µg/mice) or vehicle (PBS) every day for 30 days (**Fig. 2A**). The animals were monitored daily for any morbidity or adverse morphological changes. It was observed that ovariectomy caused significant gain in the body weight of animals as compared to the SHAM group. Interestingly, the overall weight of IL-3 treated mice remained unchanged and was closer to SHAM operated, suggesting that IL-3 may play a role in moderating the body weight increase commonly observed with post-estrogen deficiency (**Fig. 2B**). Additionally, as reported, a marked reduction in uterine weight was observed in the OVX mice due to uterine atrophy. However, IL-3 did not show any effect on this decrease in uterine weight, suggesting that IL-3 does not exhibit any estrogen-like activity on uterus (**Fig. 2C**).

**Figure 2.**
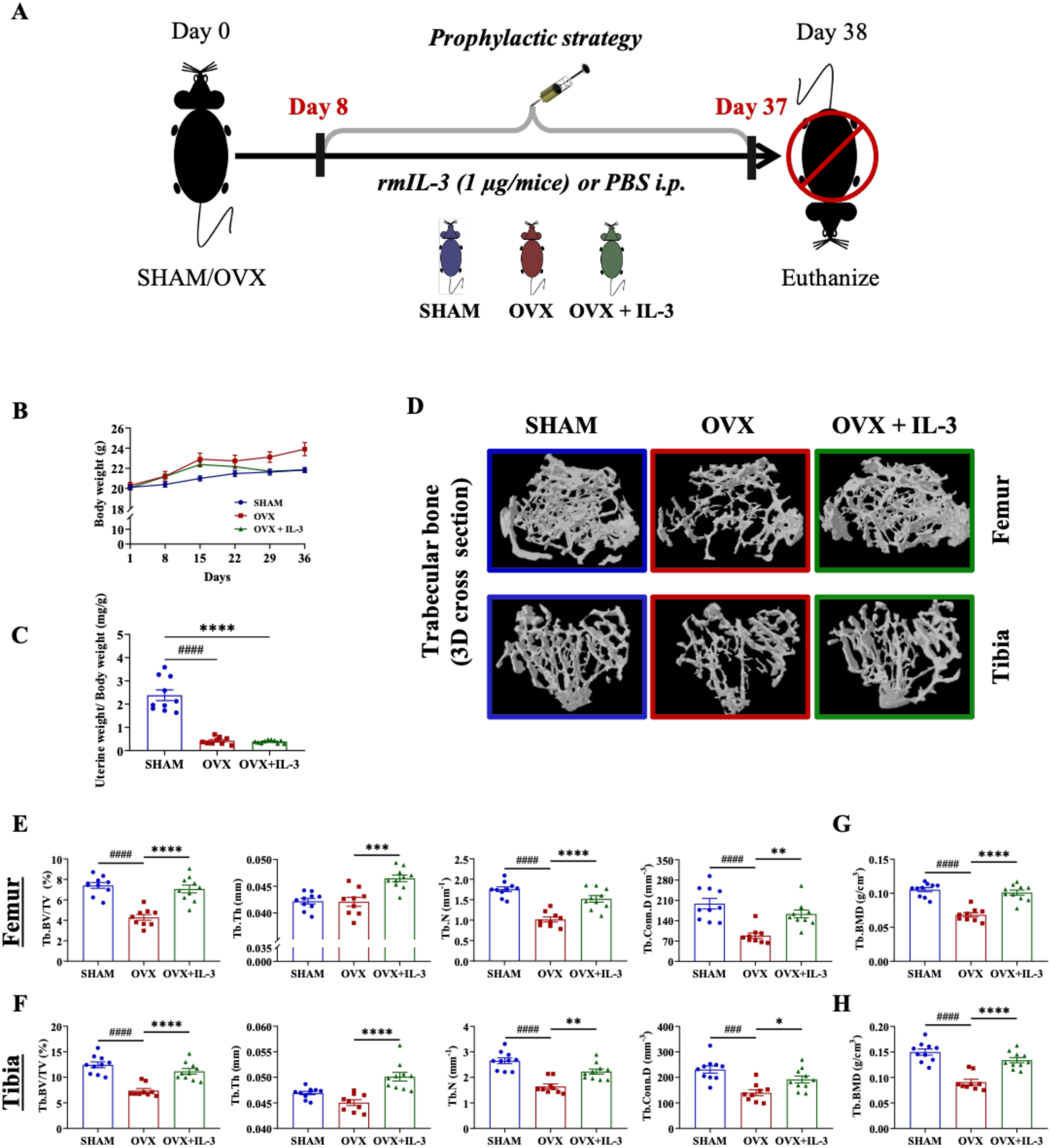
IL-3 prevents OVX-induced trabecular bone loss in OVX mice. **(A)** Prophylactic testing study involved ovariectomy of 8-10 weeks old mice, followed by i.p. administration of IL-3 (1 µg/day/mice), from day 8 post-surgery and for a duration of 30 days. **(B)** Body weight and (**C**) uterine weight were measured every week, and on day 38, mice were euthanized and bones were subjected to µ-CT. (**D)** shows representative reconstructed cross sectional 3D images of distal femur metaphyseal and proximal tibia metaphyseal trabecular bones. **(E, F)** Quantitative histomorphometry of trabecular bone indices, including BV/TV, Tb.N, and Tb.Conn.D (G, H) shows Tb.BMD. (n=9-10 mice/group, #### p ≤ 0.0001 vs. SHAM; ** p ≤ 0.01; *** p ≤ 0.001; **** p ≤ 0.0001 vs. OVX.

Ovariectomy-induced osteoporosis differently impacts the bone across various bone sites depending on the mouse strain (*15*). Therefore, in our study we quantified the bone loss in regions of distal femur and proximal tibia using μ-CT. Quantitative analysis of the histomorphometric data of distal femur demonstrated that OVX caused a dramatic loss of bone mass, as indicated by a marked decrease in BV/TV, Tb.Th, Tb.N, and Conn.D, as compared the SHAM group (**Fig. 2E**). Consequently, there was a significant increase in Tb.Sp in OVX mice when compared with SHAM. Intriguingly, IL-3 treated OVX mice were observed to retain the proximal femur trabecular bone parameters, including BV/TV, Tb.Th, Tb.N. and Conn.D. To be noted, all these trabecular parameter values were comparable to that of SHAM. Similarly, at the proximal tibia metaphysis in OVX mice, IL-3 alleviated OVX-induced trabecular bone loss parameters, including BV/TV, Tb.Th, Tb.N, and Conn.D. (**Fig. 2F**). To our surprise, IL-3-treatment prevented OVX-induced loss of bone strength as indicated by a bone mineral density (BMD) which is comparable to SHAM and dramatically higher than the vehicle-treated OVX mice at both DFM and PTM (**Fig. 2G and H**). In addition, IL-3 improved other parameters of trabecular bone at both distal femur and proximal tibia, such as Tb.Sp, Tb.Pf, and SMI **(Table S2 and S3)**. Representative qualitative μ-CT -generated 3D images underlines the bone protection action of IL-3 in ovariectomy-induced bone loss model (**Fig. 2D**). These observations were further supported by histological analysis of femur tissues where IL-3 was clearly observed to reduce the deleterious effects of ovariectomy on trabecular bone loss in osteoporotic animals (**Suppl. Fig. 1**). Taken together, these data suggest a prophylactic application of IL-3 in preventing estrogen-induced bone loss in mice.

### Therapeutic action of IL-3 causes new bone formation in OVX mice

In PMO, osteoblasts’ differentiation and bone formation functions are abrogated. Thus, there is a need for drugs that can protect osteoblasts from these negative effects of estrogen deficiency. Previously, our group reported that IL-3, in vitro, promotes osteoblast differentiation of human bone marrow-derived mesenchymal stem cells from healthy individuals (*10*). Thus, IL-3 may have a potential therapeutic implication in promoting new bone formation in treating conditions such as PMO. In the prophylactic strategy (strategy-I), it was observed that administration of IL-3 from day 8 post-OVX until day 37 effectively prevented OVX-induced trabecular bone loss, possibly by inhibiting osteoclast differentiation. However, the therapeutic bone anabolic potential of IL-3 in treating osteoporosis has not yet been explored. To address this gap, we evaluated the bone anabolic potential of IL-3 in OVX mice after development of osteoporosis, marked by establishment of significant bone loss. In the strategy-II, the animals were ovariectomized and housed for 4 weeks to develop measurable significant bone loss (day 30 post-OVX), followed by daily IL-3 administration, starting from day 30 and for an additional 4 weeks (day 59 post-OVX) (**Fig. 3A**). On day 60 post-OVX, mice were euthanized and physiological and trabecular bone parameters were studied.

**Figure 3.**
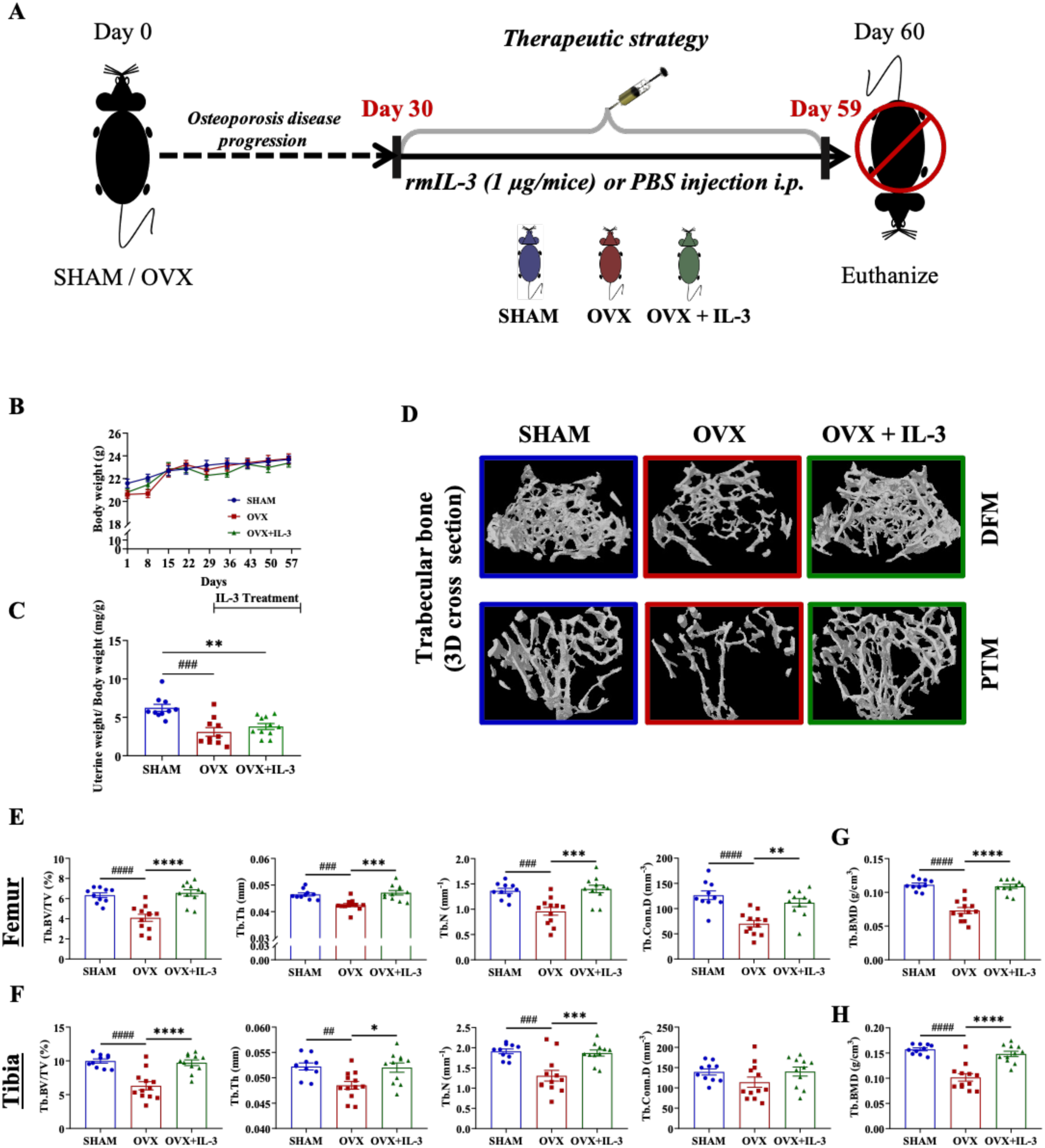
Therapeutic role of IL-3 in restoration of trabecular bone in DFM and PTM of OVX mice. **(A)** Therapeutic study involved ovariectomy of 8-10 weeks old female C57BL/6J mice, followed by administration of IL-3, 30 days post-surgery for a duration of another 30 days. **(B)** Body and **(C)** Uterine weights of mice were measured at indicated timepoints. **(D)** Representative reconstructed cross sectional 3D images of distal femur metaphyseal and proximal tibia metaphyseal bone, assessed by µ-CT, subsequent **(E, F) Histo**morphometric measurement of trabecular bone indices, including Tb.BV/TV, Tb.N, Tb.Conn.D, and (G, H) Tb.BMD, were evaluated. (n=10-12 mice/group; # p ≤ 0.05; ## p ≤ 0.01; ### p ≤ 0.001; #### p ≤ 0.0001 vs. SHAM; * p ≤ 0.05; ** p ≤ 0.01; *** p ≤ 0.001; **** p ≤ 0.0001 vs. OVX.

The body weight observation of all the experimental groups showed that on day 60 post-OVX, neither the OVX surgery nor IL-3 had any negative impact on body weight of animals (**Fig. 3B**). Furthermore, uterine/ body weight ratio was notably decreased in both OVX group and IL-3 treated-OVX group as compared to SHAM, suggesting that IL-3 doesn’t display any estrogen-like properties (**Fig. 3C**).

Further μ-CT analysis of bone, revealed a striking observation that the administration of IL-3 from day 30 to day 59 had a significant positive anabolic effect on femur as well as tibia trabecular bones. Supporting these findings, all the trabecular bone indices, including BV/TV, Tb.Th, Tb.N, and Conn.D are found to be significantly enhanced in IL-3 treated OVX mice as compared to vehicle treated OVX mice (**Fig. 3E and F**). Interestingly, femur and tibia trabecular BMD were also significantly increased in IL-3 treated mice when compared to the vehicle treated OVX group at both DFM and PTM (**Fig. 3G and H; Table S4 and S5**). μ-CT -based 3D images of distal femur and proximal tibia trabecular bone provided visual evidence that IL-3 can help osteoblasts form new bone in ovariectomy-induced bone loss model in mice (**Fig. 3D**). Based on these results, we conclude that administering IL-3 after the bone loss is established can help reverse OVX-induced bone loss by promoting new bone formation in mice.

### IL-3 has bone anabolic action on estrogen deficiency-induced cortical bone loss

In prophylactic experimental study, where IL-3 was administered before the significant bone loss was established, OVX led to a significant reduction in several cortical bone parameters, including Ct.BV/TV, Ct.Th, Ct.B.Ar/T.Ar and Ct.BMD, in both the femur and tibial diaphysis. However, the administration of IL-3 in the OVX mice showed a mild positive effect on OVX-induced cortical bone loss, as shown by a moderate improvement in the structural parameters of the cortical bone in either the femur or tibia (**Fig. 4A-C; Table S6 and S7**). Importantly, no adverse effect of IL-3 was observed on any of the cortical parameters.

**Figure 4.**
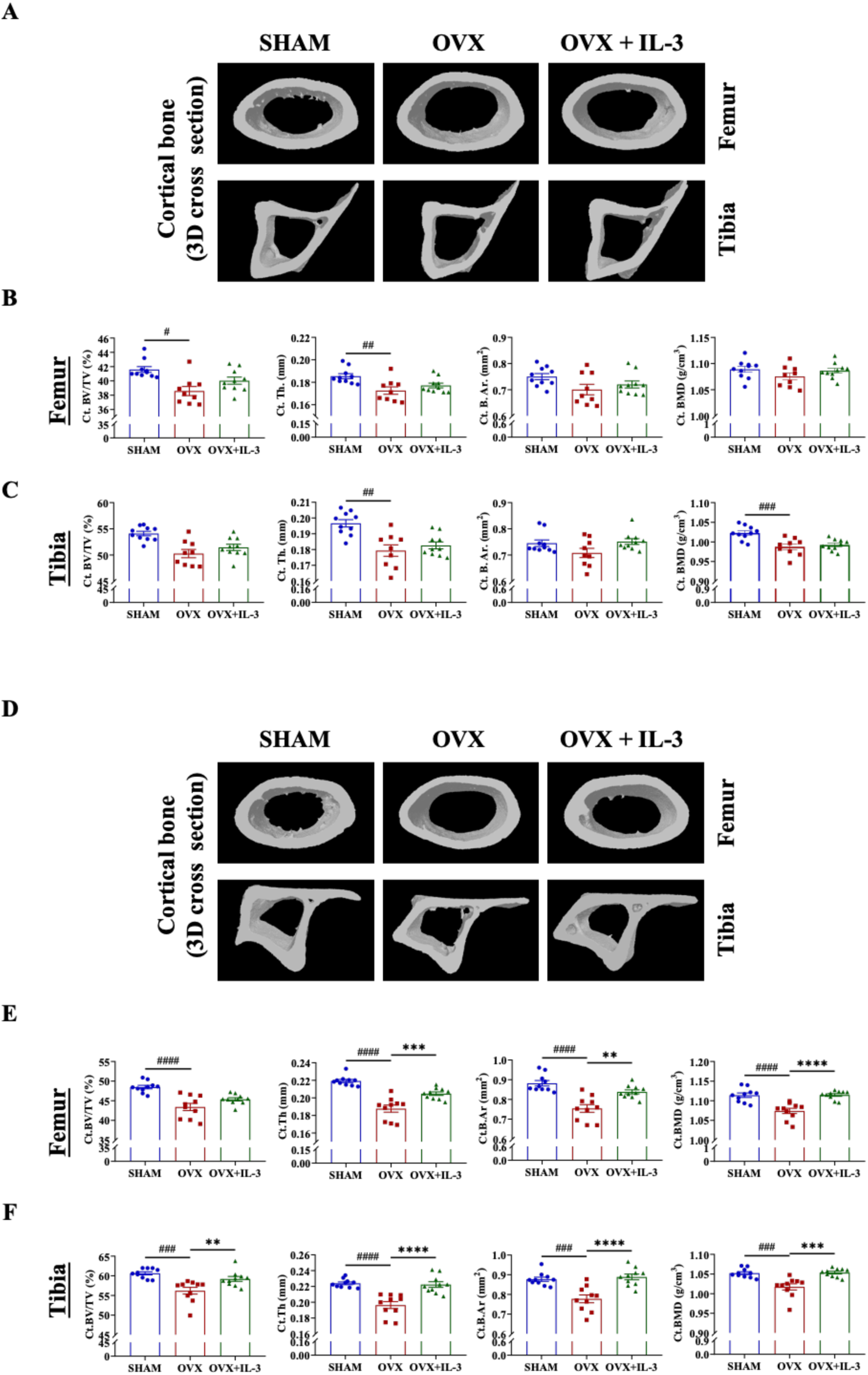
Anabolic effect of IL-3 on estrogen-deficiency-induced cortical bone loss. **(A)** Reconstructed 3D model of the cortical regions of femur and tibia from mice from prophylactic strategy. Histomorphometrical measurements of **B**, femoral and **C**, tibial cortical bone indices such as, Ct.BV/TV, Ct.Th, Ct.B.Ar, and Ct.BMD. **(D)** 3D model of the cortical regions of femur and tibia from mice from Therapeutic strategy. Histomorphometric analysis of cortical region in (**E**) femoral and (**F**) tibial bone using µ-CT. (n=9-10; # p≤ 0.05; ## p ≤ 0.01; ### p ≤ 0.001vs. OVX group)

In the therapeutic experimental strategy, two months post-OVX, the bone parameters, including Ct.BV/TV, Ct.Th, and Ct.B.Ar, showed significant reduction in cortical regions of both femur and tibia in osteoporotic mice. Interestingly, administration of IL-3 reinstated all these parameters, which were comparable to SHAM group, thus significantly improving the overall cortical microarchitecture (**Fig. 4E-F; Table S8 and S9**). Notably, the BMD was higher in IL-3 treated OVX group as compared to vehicle treated OVX group, and the BMD values were restored up to the normal range of SHAM group. All these data strongly suggest the anabolic role of IL-3 on cortical bone, in PMO condition.

### IL-3 alleviates OVX-induced increased osteoclast and osteoblast mediated bone remodeling to mediate its bone protective and bone anabolic actions with no adverse-effects in mice

Bone remodeling requires a well-coordinated balance in the actions of osteoclasts and osteoblasts to maintain bone health. In PMO, the balance in the actions of these two cells is perturbed, leading to over-activation of OC differentiation and function, and inhibition of osteoblast function, leading to a significant increase in bone loss (*15–18*). For this purpose, osteoclast- and osteoblast-specific marker gene expression in the bone tissue were analyzed by qPCR. In the prophylactic treatment strategy (**Fig. 2A**), it was observed that OVX upregulated the expression of osteoclast specific marker genes in femur and tibia, including *Ctsk, Tnfrsf11a, Mmp9, Trap, integrinβ3, Dc-stamp, and Nfatc1*, as compared to SHAM group. And, IL-3 treated group showed a trend of downregulation of all these OC marker genes mice, thus demonstrating the potential role of IL-3 in inhibiting osteoclastogenesis and bone resorption (**Fig. 5 A**). Further analysis of OB marker genes, such as *Osx, Opn, Ocn, Alp, and Col1*, in vehicle treated OVX group showed upregulation of these markers, as compared to SHAM group. However, IL-3 showed a trend of down regulation of these OB markers genes, which was comparable to SHAM group (**Fig. 5 C**). In addition, both serum CTX-I, degradation product of collagen type-1 and a marker of osteoclast function, were brought back to SHAM operated animals. A similar significant trend was seen with osteoblast marker, osteocalcin, OCN. This is suggesting that IL-3 brings back the OVX-induced accelerated bone remodeling. Intriguingly, in the therapeutic treatment strategy (**Fig. 3A**), based on the gene expression profile of OC marker genes-*Ctsk, Tnfrsf11a, Mmp9, Trap, integrinβ3, Dc-stamp, and Nfatc1*.

**Figure 5.**
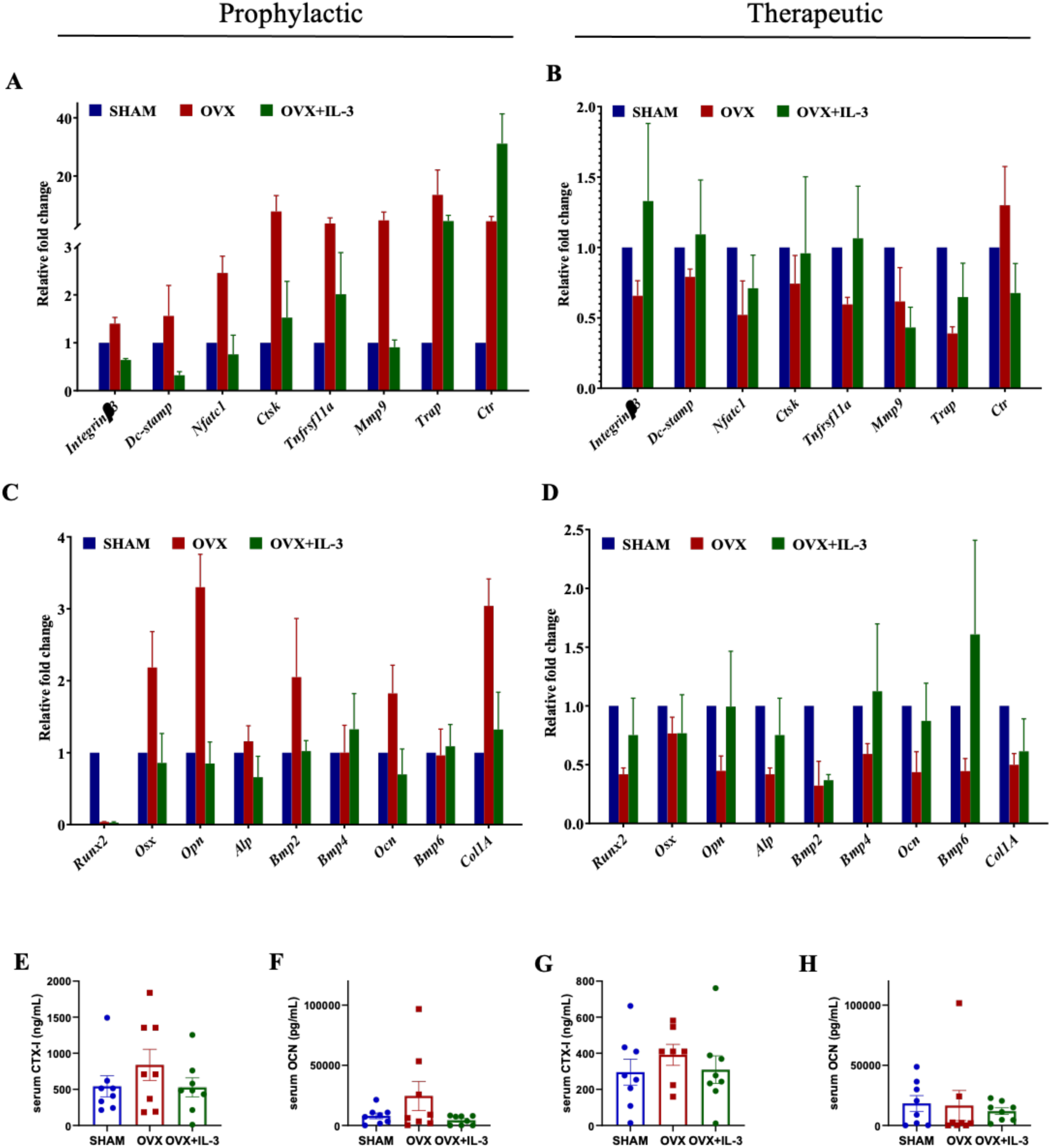
**IL-3 alters OVX-induced changes in osteoclasts- and osteoblasts-specific gene expression.** Marrow free bones from IL-3 treated animals from both Prophylactic and Therapeutic treatment strategy, as indicated, were subjected to mRNA expression profile and (**A, B**) Osteoclast-specific markers: *integrinβ3, Dc-stamp, and Nfatc1*, *Ctsk, Tnfrsf11a(RANK), Mmp9, Trap,* and *Ctr.* (B, D) Osteoblast-specific markers: *Runx2, Osx, Opn, Alp, Bmp2, Bmp4, Ocn, Bmp6,* and *Col1* were assessed by qPCR. Serum (**E, G**) CTX and (**F, H**) OCN were also measured by ELISA in both the strategies. Data is presented as mean ± SEM with n=3 mice/group for qPCR, and n=8 mice/group for ELISA.

OB marker genes-*Osx, Opn, Ocn, Alp, and Col1*, it appears that by day 60 of OVX, the bone remodeling, which was higher on day 30 post-OVX, was brought down. Interestingly, IL-3 brought the levels of expression of these genes comparable to SHAM operated group (**Fig. 5B and D**). Corroborating these findings, in IL-3 treated mice, serum c-terminal telopeptide of type-I collagen (CTX-I) levels were brought down and was comparable to SHAM group in both the prophylactic and therapeutic strategy. A similar trend was observed in the levels of serum osteocalcin (OCN) levels in IL-3 treated mice in the prophylactic treatment strategy, however, and in the therapeutic strategy, there wasn’t any difference between any of the groups of animals. Taken together, the net result of protection of bone mass we observed in the prophylactic strategy of IL-3 could be due to inhibition of OC formation and/or function. While, the bone anabolic action of IL-3 in the therapeutic strategy could be partly due to both increase OB function and inhibition of OC differentiation. Possibly, IL-3 sequentially governed the compensatory activation of bone forming osteoblasts via osteoclasts.

As observed in our study, IL-3 has a significant impact on bone tissue. We were curious whether exogenously administered caused any changes in the other vital organs. Histological examination of vital organs, including, heart, kidney, liver, and lung, revealed no apparent tissue damage attributable to IL-3 treatment when compared to both the SHAM and OVX groups of mice from prophylactic and therapeutic strategies, as observed by H&E staining (**Suppl. Fig. 4**). These findings further support the safety profile of IL-3, suggesting its potential as a therapeutic agent without adverse effects on the vital organ tissues.

### Bone protection action of IL-3 could be mediated by immunomodulation of B and T cell subsets

Various reports have emphasized on the role of estrogen deficiency in enhancement of B-lymphopoiesis (*19*). In continuation to these claims, we observed a significant augmentation of B cell subsets CD19^+^ CD45R/B220 and CD19^+^ IgM in the bone marrow, where B cells are produced and mature. OVX increased marrow and splenic B cells percentages, which is the site of B cell formation and maturation, respectively (**Fig. 6A** and **C**). Intriguingly, IL-3 treatment in OVX animals was found to effectively reduce the OVX induce B cell maturation in spleen and lymph nodes with no change in the bone marrow B cell percentages, probably indicating that IL-3 impacted the migration of B cells into secondary lymphoid organs such as in spleen and lymph nodes. Since osteoporosis is a systemic disease and IL-3 was administered intraperitoneally, B cell subsets in the peritoneal fluid were also analyzed, however neither OVX nor IL-3 injected group showed any changes in the B cell subsets (**Fig. 6D**). Estrogen deficiency leads to increase in pathogenic T cells subsets, including Th1, Th2, and Th17, and reduced immunosuppressive T cells subsets called Tregs. Also, decreased expression of CXCR4 on Treg cells and results in their reduction in bone marrow of OVX mice(*19*)To determine whether the protective effect of IL-3 treatment is solely due to its direct action on bone cells or also mediated indirectly through immune cells, such as Treg cells, we analyzed specific cell markers, including CD4 and FOXP3, in the spleen, lymph nodes, bone marrow, and peritoneal fluid of the experimental animals. IL-3 administration increased the percentage of CD4+ FOXP3+ Treg cells in OVX mice, suggesting its role in promoting Treg cell expansion in spleen, lymph node, and peritoneal fluid (**Fig. 6E, F and H**). However, no significant effect, neither decline nor enhancement was observed in any other Th cell subsets in any lymphoid compartments, neither by OVX nor upon administration of IL-3 in OVX mice (**Fig. 6E-H**), except for Th17 subsets, which were significantly downregulated in the lymph nodes of IL-3 treated OVX mice. Nonetheless, further experiments are required to elucidate the intricate details of the action of IL-3 on B and T cell biology, specifically in context of estrogen deficiency induced bone remodeling.

**Figure 6.**
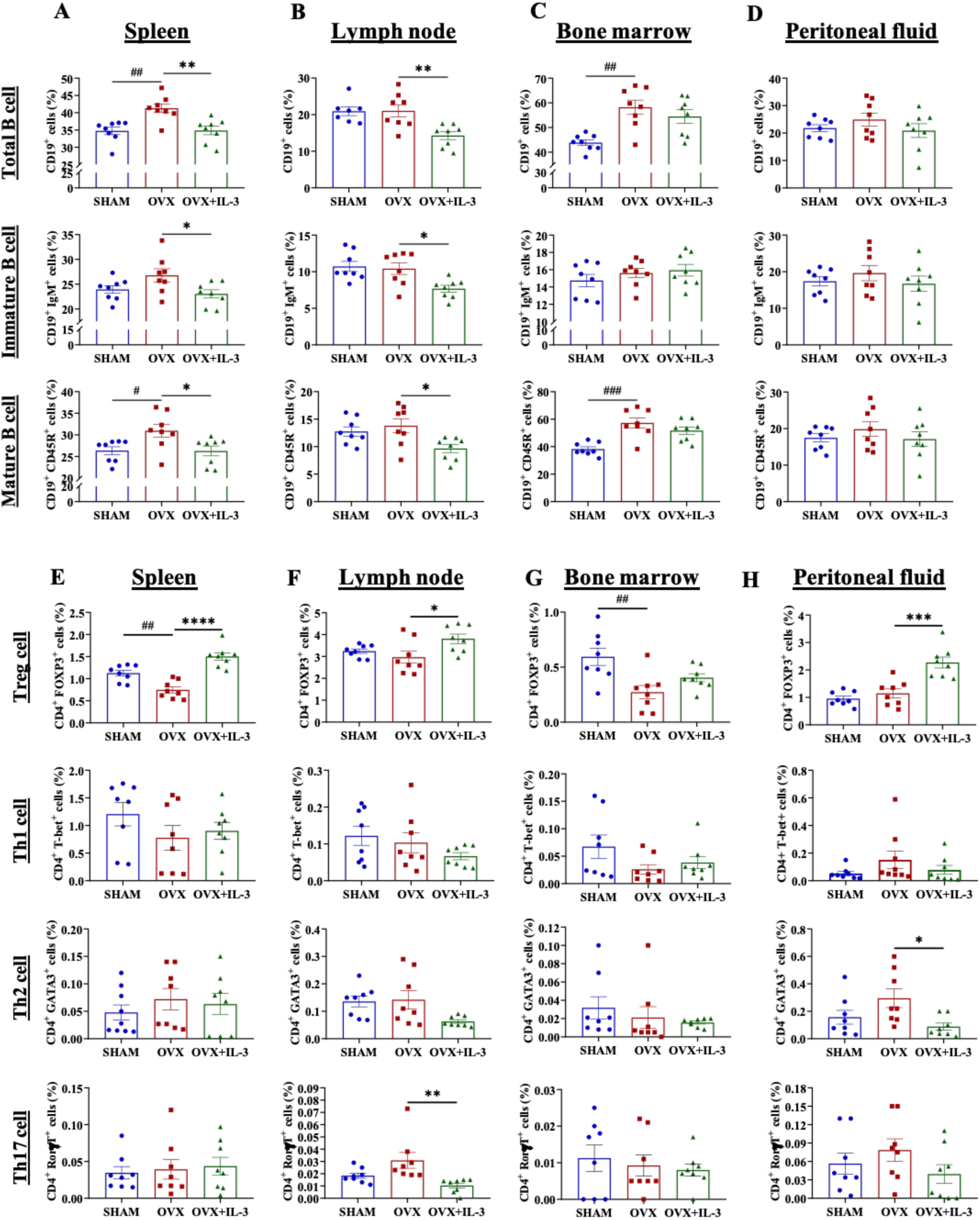
**IL-3 modulates estrogen deficiency-induced pathogenic B and T cell subsets.** Immune-phenotyping of both B and Th cell subsets from prophylactic strategy mice was carried out on spleen (**A, E**), lymph node (**B, F**), bone marrow (**C, G**), and peritoneal fluid (**D, H**), using flow cytometry. (n=7-8 mice/ group; **p < 0.01; #p ≤ 0.05, ####p < 0.0001 vs. OVX group).

Since IL-3 was discovered as a haematopoiesis factor, and is a critical factor affecting all the blood cells of both myeloid and lymphoid origin. Therefore, we assessed the haematological parameters in all groups of mice from both prophylactic and therapeutic strategies. It appears that administration of IL-3 of 1ug/day/mice for 30 days did not cause any detrimental changes in haematological parameters, including haemoglobin, haematocrit, white blood cells, and platelet count (**Table S10 and S11**). Thus, suggesting that IL-3 has the potential to be used as a drug in treating PMO with no major toxic side effects.

### IL-3^-/-^ mice display osteopenic phenotype with both less bone mass and mineral density

The exogenously administered recombinant mouse IL-3 in OVX mice protected from bone loss and also helped neo-bone formation. We were curious if endogenous IL-3 has any role in the physiology of mice, including the bone health. Thus, we utilized IL-3 KO mice and determined their basal bone microarchitecture indices using µ-CT analysis.

The body weight measurement revealed that IL-3 KO mice (**Fig. 7A and H**), both male and female, were heavier as compare to their WT counterparts (**Fig. 7B and I**). As IL-3 is known to regulate haematopoiesis and related immune responses, a detailed analysis of haematological parameters revealed no major differences, except that the % neutrophils were higher in male IL-3^-/-^ as compared to WT counterparts (**Table S12**). Interestingly, a thorough investigation of biochemical indices in the plasma samples displayed lower levels of ALP in the IL-3^-/-^ in both males and females, a lower blood cholesterol in the IL-3^-/-^ males as compared to their WT counterparts (**Table S13**).

**Figure 7.**
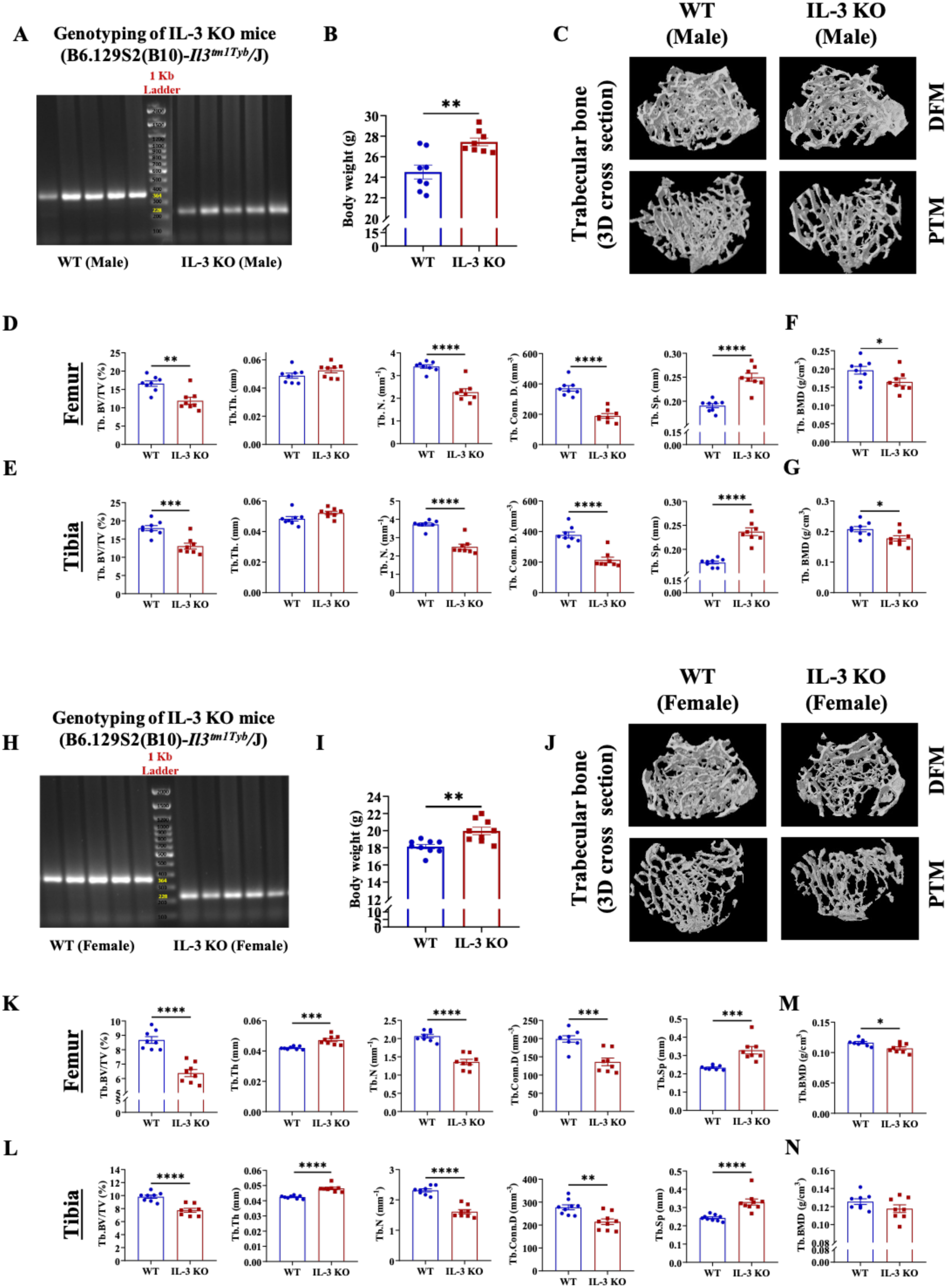
IL-3^-/-^ mice display osteopenia and is sex-independent. **A**, Male and **H**, Female IL-3^-/-^ and WT littermates were genotyped. Body weight of 8-10 week old IL-3^-/-^ and WT littermates of **B**, male and **I**, female were examined , and their respective reconstructed 3D micro-CT images (**C**, **J**) of trabecular bones and quantitative histomorphometric measurements of trabecular bone indices such as BV/TV, Tb.N, Tb.Th, Conn.D, and BMD at DFM (**D**, **F and F**, **M**) and PTM (**E**, **G - L**, **N)**, were measured. (n=8 mice/ group; * p ≤ 0.05; ** p ≤ 0.01; *** p ≤ 0.001; **** p ≤ 0.0001 vs. WT littermates.

Further bone phenotyping, using µ-CT, intriguingly, 3D reconstructed images of IL-3^-/-^ mice revealed significant lesser trabecular bone in both male and female mice (**Fig. 7C and J**). Quantitively, IL-3^-/-^ displayed a dramatic lower value for BV/TV, Tb.N, Conn.D, and with a corresponding increase in Tb.Sp, in both DFM and PTM. However, for unknown reasons, Tb.Th did not show any change in IL-3^-/-^ mice. Interestingly, IL-3^-/-^ exhibited poor bone strength as indicated by a significant reduction in trabecular BMD at both DFM (**Fig. 7F and M**) and PTM **(Fig.7G and N).** In summary, IL-3 plays a significant role in regulating in bone homeostasis, with no major roles in adult haematopoiesis, where its role has been considered important.

### IL-3^-/-^ mice are not resistant to OVX-induced bone loss

In our study, exogenously administered recombinant IL-3 had dual actions of both protecting bone from OVX-induced loss and forming new bone. In addition, IL-3^-/-^ mice exhibited poor bone mass and density. Thus, we were curious to know if ovariectomizing IL-3^-/-^ mice will make the bone phenotype worse. Body weight measurement during the course of bone loss post-OVX and until day 30 (**Fig. 8A**), suggested that IL-3^-/-^ mice gained more weight than the WT counterparts (**Fig. 8B**). Confirming our previous result of bone phenotype in IL-3^-/-^, it is clear that SHAM operated IL-3^-/-^ mice have lower bone mass than WT counterparts at both DFM and PTM. Upon OVX of IL-3^-/-^ there was a significant bone loss in both DFM and PTM as indicated by significantly reduced BV/TV, Tb.N, Conn.D, and consequently, increased Tb.Sp. (**Fig. 8F and G; Table S14 and S15**). The reconstructed µ-CT images shown depicted the bone loss induced by OVX in IL-3^-/-^ and WT (**Fig. 8D and E**). Moreover, BMD has also declined in the IL-3^-/-^ upon OVX at both femur and tibial bones (**Fig. 8H and I**). All these data, suggested that IL-3 is essential in maintaining trabecular bone mass and strength, and that bone the estrogen-deficiency induced bone loss or maintenance of bone mass by estrogen is independent of IL-3. However, IL-3 independently regulates bone homeostasis.

**Figure 8.**
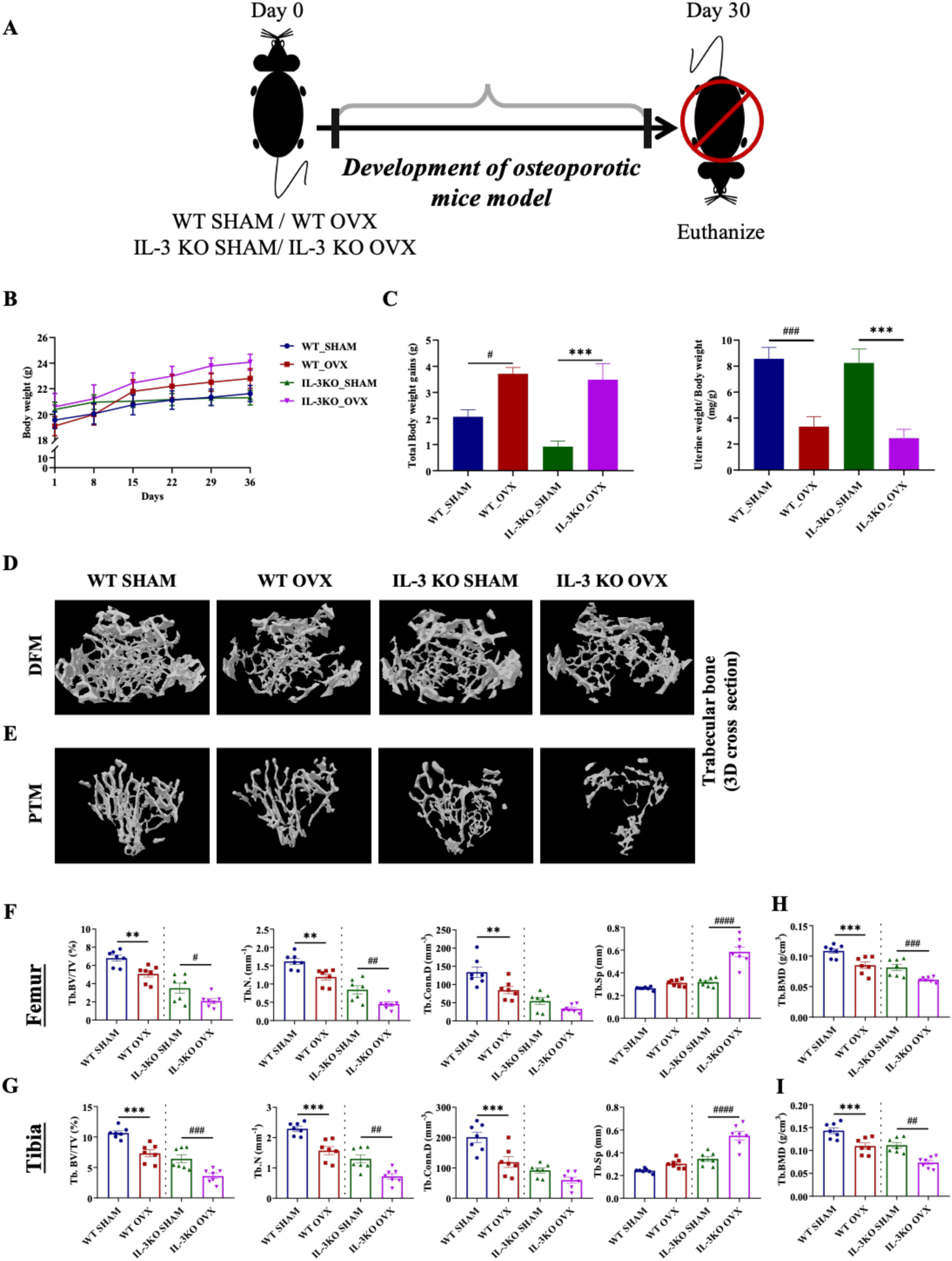
IL-3^-/-^ mice display bone OVX-like bone phenotype and are not resistant to OVX-induced bone loss. **A**, 8–10-week-old IL-3^-/-^ amd their WT littermates were ovariectomized and (**B**) body weight (**C**) uterine weight was measured at the indicated time points until day 30. Post-euthanasia bones were subjected to µ-CT. 3D reconstructed micro-CT images of trabecular bone at (**D)** DFM and (**E)** PTM. Further, quantitative histomorphometric measurements of trabecular bone indices such as BMD, BV/TV, Tb.N, Tb, Th, Conn.D, and BMD is shown for both (**F**, **H**) DFM and (**G**, **I**) PTM. * p ≤ 0.05; ** p ≤ 0.01; *** p ≤ 0.001; **** p ≤ 0.0001 vs. WT littermates.

## Discussion

Bone remodeling is the well-coordinated process of bone resorption by osteoclasts and new bone formation by osteoblasts for maintenance of bone homeostasis. Disruptions of this cycle result in bone diseases such as osteoporosis, osteopetrosis, rheumatoid arthritis, Paget’s disease, and bone malignancies. PMO is one such condition associated with estrogen deficiency, marked by low mass and bone mineral density causing skeletal fragility, eventually leading to significant bone loss and increased risk of fractures. Clinical management of PMO includes an array of drugs which are mostly anti-resorptive and the only bone anabolic drug available are the synthetic analogs of parathyroid hormone and Romosozumab. However, both of these anabolic agents are used for patients with higher risk of fractures are associated with side serious effects when used long-term (*20*). This emphasises the need of novel bone anabolic agents with no or negligible side effects. Thus, we investigated the role of interleukin-3 as a novel therapeutic agent in treating PMO.

To thoroughly examine the role of IL-3 at different stages of the PMO development and prognosis, we applied two different strategies in a mouse model of ovariectomy-induced bone loss. In our first strategy, which we called prophylactic strategy, where IL-3 was administered 7 days post-ovariectomy for a duration of 30 days, we observed that the IL-3 injected OVX mice displayed a comparable trabecular bone parameters with SHAM, at both distal femur and proximal tibia. Moreover, it was fascinating to see that IL-3 treatment protected the mice from OVX-induced loss of bone mineral density (BMD). In our second strategy, called as therapeutic strategy, where IL-3 was administered 30 days post-ovariectomy for a duration of additional 30 days, we did observe significant cortical bone loss in addition to trabecular bone loss in the OVX mice. Intriguingly, IL-3 restored the lost trabecular and cortical bone parameters at both distal femur and proximal tibia. Also, the bone micro-architecture of IL-3 treated mice was similar to SHAM operated group. Our results from both the aforementioned strategies led us to conclude that IL-3 played a significant role in bone remodeling, and not only did IL-3 prevented the OVX-induced bone loss but also had a therapeutic bone anabolic action. To further substantiate the role of IL-3 in the maintenance of bone homeostasis, we examined the bone microarchitecture of IL-3^-/-^ mice. The μ-CT analysis revealed that the basal level of trabecular bone indices in IL-3^-/-^ mice were equivalent to osteoporotic mice, suggesting that depletion of IL-3 resulted in osteoporotic bone phenotype in sex-independent manner. This confirms our previous speculation, where administration of IL-3 was able to prevent bone loss and also efficiently repaired bone microarchitecture in osteoporotic mice. qPCR based studies and serum provides further evidence that IL-3 acted on both OC and OB and brought changes in their marker genes in a manner that was opposite to the trend seen in OVX mice, suggesting IL-3 acted to establish the homeostatic bone remodelling as the marker expression was similar to the expression pattern in SHAM operated mice. A decrease in serum level of CTX-I, a marker of osteoclast function, further confirmed that IL-3 is anti-osteoclastogenic in vivo. This was evident in the prophylactic and therapeutic strategies. However, OCN levels, a marker of OB function, increased in the OVX mice was brought down to SHAM levels upon IL-3 treatment. However, in the therapeutic strategy, for unknown reason, the serum OCN levels measured were all the same in all the groups. This is surprising, as the new bone formation observed with μ-CT is robust. However, there is a need for more studies required to prove that the new bone formation is indeed via IL-3’s anabolic action on osteoblasts.

PMO, is an estrogen deficiency-induced inflammatory disorder. To be more precise, estrogen deficiency causes production of pathogenic B and T cells, which secrete inflammatory molecules, including TNF-α, IL-17, IL-1, IL-6 (*21–25*). These molecules are reported to be pro-OC and anti-OB, resulting in massive bone loss (*1, 21, 26–29*). We have some knowledge from our previous studies on the immunomodulatory role of IL-3 in vitro in that it enhances Treg cell differentiation and inhibits Th17 subset differentiation. However, the role of IL-3 in T cell and B cell subset differentiation in PMO is limited. Thus, we investigated the role of IL-3 in modulating the action of these B cell and T cell subsets in our OVX mice model. It was observed that IL-3 acted on the lymphoid component of the hematopoietic system, and caused a reduction in the formation of Th17 cells, and conversely, IL-3 increased the percentages of Tregs. Both these results supported the immunomodulatory role of IL-3, further suggesting that IL-3 treatment can efficiently safeguard the overall bone health by minimising the ovariectomy-induced inflammatory condition in osteoporotic mice. However, we have not performed cytokine array to study the inflammation status of these mice. Thus, the immuno-modulatory actin of IL-3 and its direct action on both OC and OB partly explains the action of IL-3 in the maintenance and treatment of OVX-induced bone loss.

At molecular level, we have previously reported in vitro in mice, humans, and rats, that IL-3 inhibits OC differentiation induced by either RANKL or TNF-α and/or other pro-inflammatory cytokines, including IL-1 and IL-6 and TGFs (*7, 9*). Mechanistically, IL-3 inhibits RANKL-induced NF-kB pathway and TNF-α-induced-c-fos pathway (*5, 30*). Moreover, IL-3 enhances osteogenic differentiation of human bone marrow-derived mesenchymal stem cells (*10*). Interestingly, IL-3 protects the osteoblasts from the negative action of TNF-α on osteoblast differentiation (*11*). Further, IL-3 decreases the osteoblast expression of secretory form of RANKL via down-regulation of MMPs in mouse (*31*). Our in vivo findings where administration of IL-3 in osteoarthritis (OA) and rheumatoid arthritis (RA) mouse models not only protected the cartilage and bone from inflammation-induced damage, but also prevented the onset of osteoarthritis and inflammatory conditions, thus suggesting the involvement of IL-3 in immunomodulation as well as bone protection (*6, 12, 32*).

Clinically proven drugs in the management of osteoporosis includes, bisphosphonates, which are anti-resorptive-which are known to cause osteonecrosis of jaw if used longer, and anabolic agents like Teriparatide increases osteosarcoma risk, while Denosumab causes hypocalcaemia, unusual hip fractures and osteonecrosis. Romosozumab, a new anabolic drug, is an antibody against sclerostin, which inhibits bone formation, and is associated with negative impact on cardiovascular health (*33*). Thus, each of these drugs require careful consideration when in use. Thus, it was necessary to evaluate if IL-3 has any toxic side effects in our OVX mice. We found no toxic effects of IL-3 on vital organs, including heart, liver, and kidney by histology. Since, IL-3 is a growth factor for blood cells. We found no major changes in the complete blood count (CBC) profile of the IL-3 administered OVX mice. Previous clinical studies in lung cancer patients undergoing chemotherapy in 90’s showed that IL-3 is well-tolerated in humans up to a dose of 16ug/kg/day, and headache was considered to be dose limiting (*34*). However, IL-3^-/-^ mice did show higher percentages of neutrophils only in males. The serum biochemistry reveals dramatic decline in ALP in both male and female IL-3^-/-^ mice and male sex specific decline in total cholesterol in IL-3^-/-^ mice. These suggest that IL-3 may have more success in the clinic in the treatment of PMO with mild to no side effects. In summary, our previous reports and the current findings in PMO strongly suggest that IL-3 has the potential to be a novel drug due to several advantages-a promising inhibitor of bone resorption, promoter of bone formation, and an immunomodulatory agent, and most importantly may exhibit less detrimental side effects. However, a thorough investigation is required to prove some of aspects which the current study could not cover, including the dynamic histomorphometric analysis to show new bone formation, and any in vivo mechanistic studies in the OVX model to identify the actual molecular targets, which could open a new avenue of research. Moreover, we have not tested IL-3 in aging-induced bone loss, which could be addressed in our subsequent studies.

### Ethics approval and consent to participate

This study protocol involving the use of animals received approval from the Institutional Animal Ethics Committee (Project no. IAEC/2016/B-277).

## Availability of data and materials

The datasets generated and/or analyzed during the current study are available from the corresponding author upon reasonable request

## Authors’ contributions

V.P. and M.R.W. conceptualized and designed the methods; V.P., S.B., J.K., G.P., A.P., K.E. performed the experiments; V.P. and M.R.W. analyzed the data, V.P. and S.B. prepared the figures, and wrote the manuscript. All authors reviewed the manuscript and approved for submission.

## Acknowledgements

We dedicate this research paper in the loving memory of our mentor, late Dr. Mohan R. Wani, who unfortunately passed away on 14th April, 2024. This work was initiated under the guidance of the late Dr. Mohan R. Wani, who unfortunately passed away on 14^th^ April 2024. We gratefully acknowledge his contributions and mentorship. We thank all the central facilities of NCCS and their staff for the support during this work.

## Bibliography

1. M. N. Weitzmann, R. Pacifici, Estrogen deficiency and bone loss: an inflammatory tale. J Clin Invest 116, 1186–1194 (2006).

2. S. P. Tuck, R. M. Francis, Osteoporosis. Postgrad Med J 78, 526 (2002).

3. J. T. Lin, J. M. Lane, Nonpharmacologic management of osteoporosis to minimize fracture risk. Nat Clin Pract Rheumatol 4, 20–25 (2008).

4. A. Y. Parra-Torres, M. Valdés-Flores, L. O. and R. Velázquez-Cruz, A. Y. Parra-Torres, M. Valdés-Flores, L. O. and R. Velázquez-Cruz, Molecular Aspects of Bone Remodeling. Topics in Osteoporosis (2013), doi:10.5772/54905.

5. S. M. Khapli, L. S. Mangashetti, S. D. Yogesha, M. R. Wani, IL-3 acts directly on osteoclast precursors and irreversibly inhibits receptor activator of NF-kappa B ligand-induced osteoclast differentiation by diverting the cells to macrophage lineage. J Immunol 171, 142–151 (2003).

6. S. D. Yogesha, S. M. Khapli, R. K. Srivastava, L. S. Mangashetti, S. T. Pote, G. C. Mishra, M. R. Wani. IL-3 inhibits TNF-alpha-induced bone resorption and prevents inflammatory arthritis. J Immunol. 2009 Jan 1;182(1):361–70. doi: 10.4049/jimmunol.182.1.361.

7. S. D. Yogesha, S. M. Khapli, M. R. Wani, Interleukin-3 and granulocyte-macrophage colony-stimulating factor inhibits tumor necrosis factor (TNF)-α-induced osteoclast differentiation by down-regulation of expression of TNF receptors 1 and 2. Journal of Biological Chemistry 280, 11759–11769 (2005).

8. V. Piprode, K. Singh, A. Kumar, S. R. Joshi, M. R. Wani, IL-3 inhibits rat osteoclast differentiation induced by TNF-a and other pro-osteoclastogenic cytokines. J Biosci (Rajshari) 46 (2021), doi:10.1007/s12038-021-00181-3.

9. N. Gupta, A. P. Barhanpurkar, G. B. Tomar, R. K. Srivastava, S. Kour, S. T. Pote, G. C. Mishra, M. R. Wani, IL-3 Inhibits Human Osteoclastogenesis and Bone Resorption through Downregulation of c-Fms and Diverts the Cells to Dendritic Cell Lineage. The Journal of Immunology 185, 2261–2272 (2010).

10. A. P. Barhanpurkar, N. Gupta, R. K. Srivastava, G. B. Tomar, S. P. Naik, S. R. Joshi, S. T. Pote, G. C. Mishra, M. R. Wani, IL-3 promotes osteoblast differentiation and bone formation in human mesenchymal stem cells. Biochem Biophys Res Commun 418, 669–675 (2012).

11. V. Piprode, S. Behera, J. Karhade, S. T. Mhaske, M. R. Wani, Priming of mouse pre- osteoblasts with Interleukin-3 attenuates TNF-α-induced-inhibition of osteoblast differentiation. BMC Musculoskelet Disord 26, 663 (2025).

12. S. Kour, M. G. Garimella, D. A. Shiroor, S. T. Mhaske, S. R. Joshi, K. Singh, S. Pal, M. Mittal, H. B. Krishnan, N. Chattopadhyay, A. H. Ulemale, M. R. Wani, IL-3 Decreases Cartilage Degeneration by Downregulating Matrix Metalloproteinases and Reduces Joint Destruction in Osteoarthritic Mice. The Journal of Immunology 196, 5024–5035 (2016).

13. A. Sophocleous, A. I. Idris, Rodent models of osteoporosis. Bonekey Rep 3, 614 (2014).

14. S. M. Khapli, G. B. Tomar, A. P. Barhanpurkar, N. Gupta, S. D. Yogesha, S. T. Pote, M. R. Wani, Irreversible inhibition of RANK expression as a possible mechanism for IL-3 inhibition of RANKL-induced osteoclastogenesis. Biochem Biophys Res Commun 399, 688– 693 (2010).

15. M. L. Bouxsein, K. S. Myers, K. L. Shultz, L. R. Donahue, C. J. Rosen, W. G. Beamer, Ovariectomy-Induced Bone Loss Varies Among Inbred Strains of Mice. Journal of Bone and Mineral Research 20, 1085–1092 (2005).

16. P. Garnero, E. Sornay-Rendu, M. C. Chapuy, P. D. Delmas, Increased bone turnover in late postmenopausal women is a major determinant of osteoporosis. J Bone Miner Res 11, 337–349 (1996).

17. B. Liang, G. Burley, S. Lin, Y. C. Shi, Osteoporosis pathogenesis and treatment: existing and emerging avenues. Cellular & Molecular Biology Letters 2022 27:*1* **27**, 1–26 (2022).

18. J. Xu, L. Yu, F. Liu, L. Wan, Z. Deng, The effect of cytokines on osteoblasts and osteoclasts in bone remodeling in osteoporosis: a review. Front Immunol 14, 1222129 (2023).

19. T. Masuzawa, C. Miyaura, Y. Onoe, K. Kusano, H. Ohta, S. Nozawa, T. Suda, Estrogen deficiency stimulates B lymphopoiesis in mouse bone marrow. J Clin Invest 94, 1090–1097 (1994).

20. D. Wu, L. Li, Z. Wen, G. Wang, Romosozumab in osteoporosis: yesterday, today and tomorrow. J Transl Med 21, 1–13 (2023).

21. R. Pacifici, Estrogen, cytokines, and pathogenesis of postmenopausal osteoporosis. J Bone Miner Res 11, 1043–1051 (1996).

22. A. M. Tyagi, K. Srivastava, M. N. Mansoori, R. Trivedi, N. Chattopadhyay, D. Singh, Estrogen deficiency induces the differentiation of IL-17 secreting Th17 cells: a new candidate in the pathogenesis of osteoporosis. PLoS One 7 (2012), doi:10.1371/JOURNAL.PONE.0044552.

23. S. Cenci, M. N. Weitzmann, C. Roggia, N. Namba, D. Novack, J. Woodring, R. Pacifici, Estrogen deficiency induces bone loss by enhancing T-cell production of TNF-alpha. J Clin Invest 106, 1229–1237 (2000).

24. C. Roggia, Y. Gao, S. Cenci, M. N. Weitzmann, G. Toraldo, G. Isaia, R. Pacifici, Up-regulation of TNF-producing T cells in the bone marrow: A key mechanism by which estrogen deficiency induces bone loss in vivo. Proc Natl Acad Sci U S A 98, 13960 (2001).

25. S. Cenci, G. Toraldo, M. N. Weitzmann, C. Roggia, Y. Gao, W. P. Qian, O. Sierra, R. Pacifici, Estrogen deficiency induces bone loss by increasing T cell proliferation and lifespan through IFN-γ-induced class II transactivator. Proc Natl Acad Sci U S A 100, 10405–10410 (2003).

26. R. Pacifici, T cells: Critical bone regulators in health and disease. Bone 47, 461–471 (2010).

27. R. Pacifici, Cytokines, estrogen, and postmenopausal osteoporosis--the second decade. Endocrinology 139, 2659–2661 (1998).

28. L. Ginaldi, M. C. Di Benedetto, M. De Martinis, Osteoporosis, inflammation and ageing. Immunity and Ageing 2 (2005), doi:10.1186/1742-4933-2-14.

29. D. S. Amarasekara, H. Yun, S. Kim, N. Lee, H. Kim, J. Rho, Regulation of Osteoclast Differentiation by Cytokine Networks. Immune Netw 18, e8 (2018).

30. J. Oh, M. S. Lee, J. T. Yeon, S. W. Choi, H. S. Kim, H. Shim, S. Y. Lee, B. S. Youn, Y. Yokota, J. H. Kim, H. B. Kwak, Inhibitory regulation of osteoclast differentiation by interleukin-3 via regulation of c-Fos and Id protein expression. J Cell Physiol 227, 1851–1860 (2012).

31. K. Singh, V. Piprode, S. T. Mhaske, A. Barhanpurkar-Naik, M. R. Wani, IL-3 Differentially Regulates Membrane and Soluble RANKL in Osteoblasts through Metalloproteases and the JAK2/STAT5 Pathway and Improves the RANKL/OPG Ratio in Adult Mice. J Immunol 200, 595–606 (2018).

32. R. K. Srivastava, G. B. Tomar, A. P. Barhanpurkar, N. Gupta, S. T. Pote, G. C. Mishra, M. R. Wani, IL-3 Attenuates Collagen-Induced Arthritis by Modulating the Development of Foxp3+ Regulatory T Cells. The Journal of Immunology 186, 2262–2272 (2011).

33. R. M. Y. Wong, P. Y. Wong, C. Liu, H. Y. Wong, M. K. Fong, N. Zhang, W. H. Cheung, S. W. Law, Treatment effects, adverse outcomes and cardiovascular safety of romosozumab – Existing worldwide data: A systematic review and meta-analysis. J Orthop Translat 48, 107 (2024).

34. P. E. Postmus, J. A. Gietema, O. Domsma, B. Biesma, P. C. Limburg, E. Vellenga, E. G. E. De Vries, Effects of recombinant human interleukin-3 in patients with relapsed small-cell lung cancer treated with chemotherapy: A dose-finding study. Journal of Clinical Oncology 10, 1131–1140 (1992).

