## Supplementary material for "Interleukin-3 as a Potential Bone Anabolic Agent in treating Postmenopausal Osteoporosis": Supple Figures and Tables

### Co-second authors

**Abstract**

Postmenopausal osteoporosis (PMO), a silent disorder caused due to estrogen deficiency, is characterized by loss of bone mass and low bone mineral density. Despite advancements in treatment, current medications often fail to fully restore bone integrity and prevent fragility fractures in the osteoporotic individuals. Thus, there is a high demand of novel therapeutic modalities in dealing with osteoporosis. Interleukin-3 (IL-3), a cytokine secreted by activated T cells, emerges as a promising therapeutic candidate. The dual role of IL-3 in inhibiting osteoclast differentiation and promoting osteoblast differentiation, offers protection against bone and joint degeneration in arthritic mice. However, its role in osteoporosis is not yet delineated. Therefore, our investigation focuses on elucidating the role of IL-3 in PMO, employing both prophylactic and therapeutic strategies in ovariectomized mice, mimicking human PMO. In our study, at the onset of osteoporosis, IL-3 treatment effectively preserved trabecular bone architecture and enhanced bone mineral density in OVX mice, particularly notable in therapeutic interventions where fully developed osteoporosis was addressed. Notably, in preventive measures, IL-3 inhibited osteoclast differentiation, thereby suppressing bone resorption, while in therapeutic approaches, it enhanced osteoblast differentiation, promoting bone formation. Irrespective of gender specific, microCT analysis of IL-$3^{-/-}$ (KO) mice showed reduced trabecular bone development compared to its respective wild type (WT) mice, highlighting essential role of IL-3 in skeletal integrity. Moreover, IL-3 increased Treg cells population while inhibiting B cell lymphopoiesis in ovariectomized mice with no adverse effects on hematopoiesis or vital organs. In conclusion, our findings collectively underscore the potential of IL-3 as a novel therapy for PMO, offering insights into its mechanisms of action and clinical applications.

**Supplementary Figure 1**

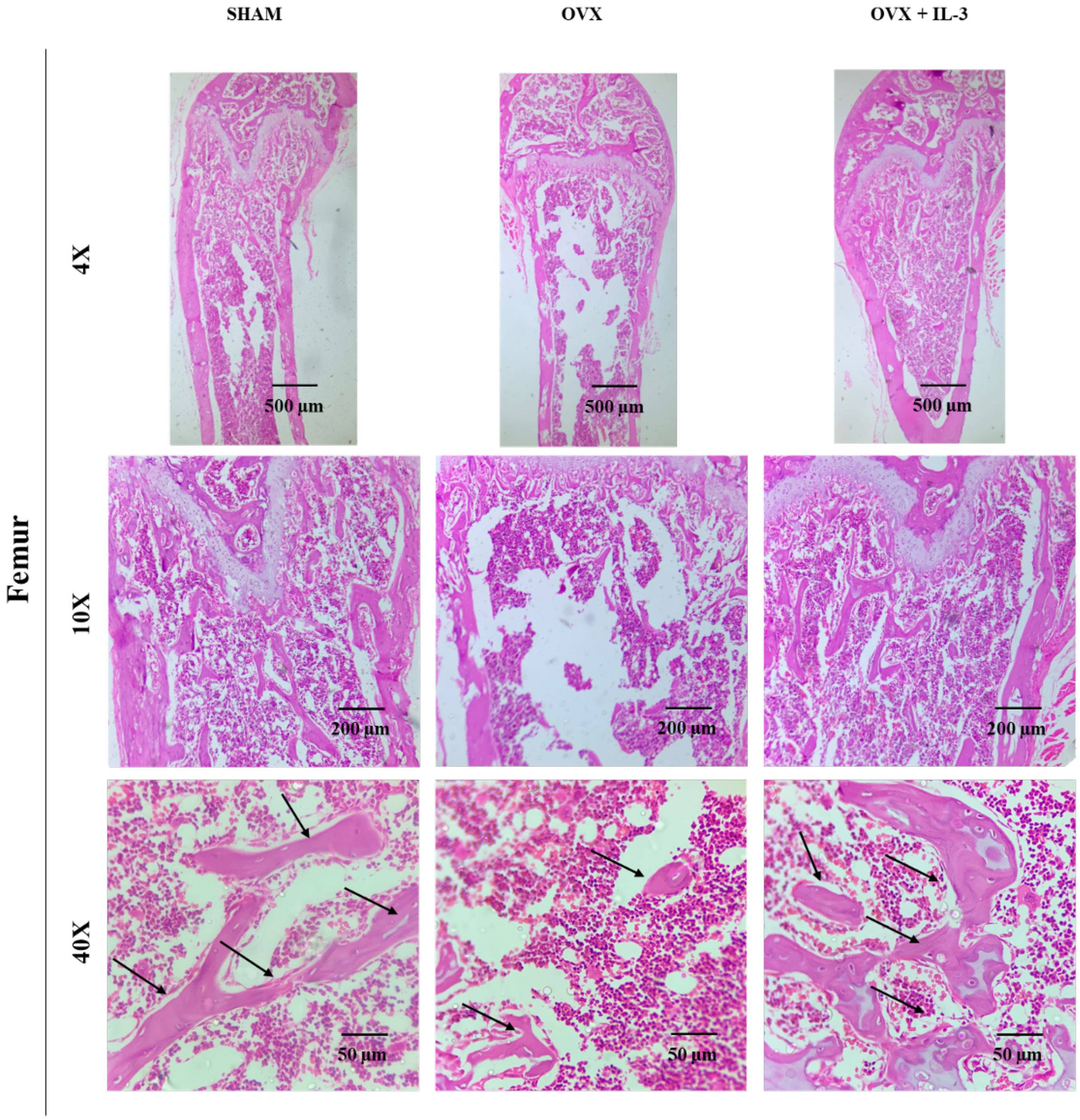

#### Supplementary Figure 2

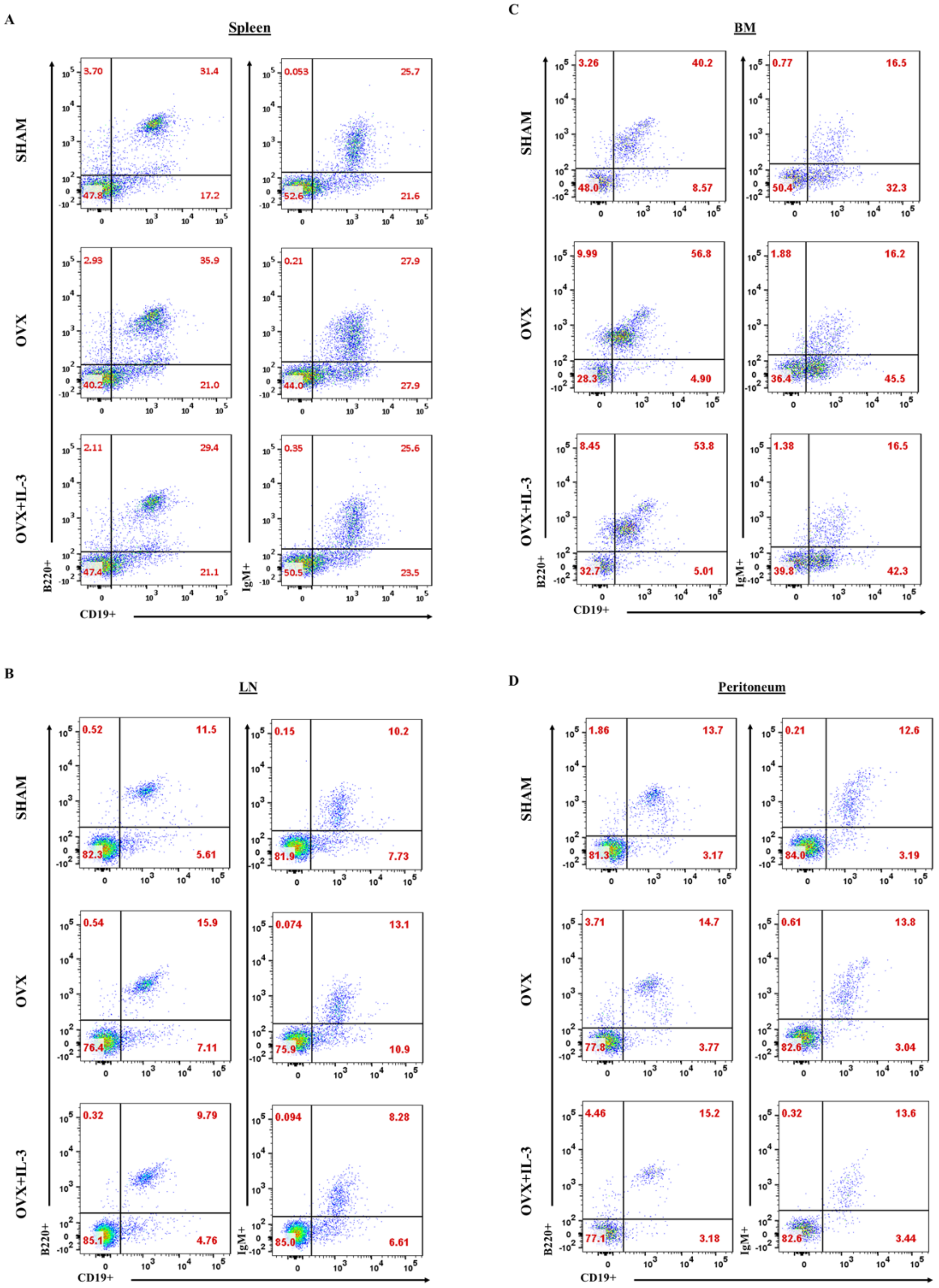

#### Supplementary Figure 3

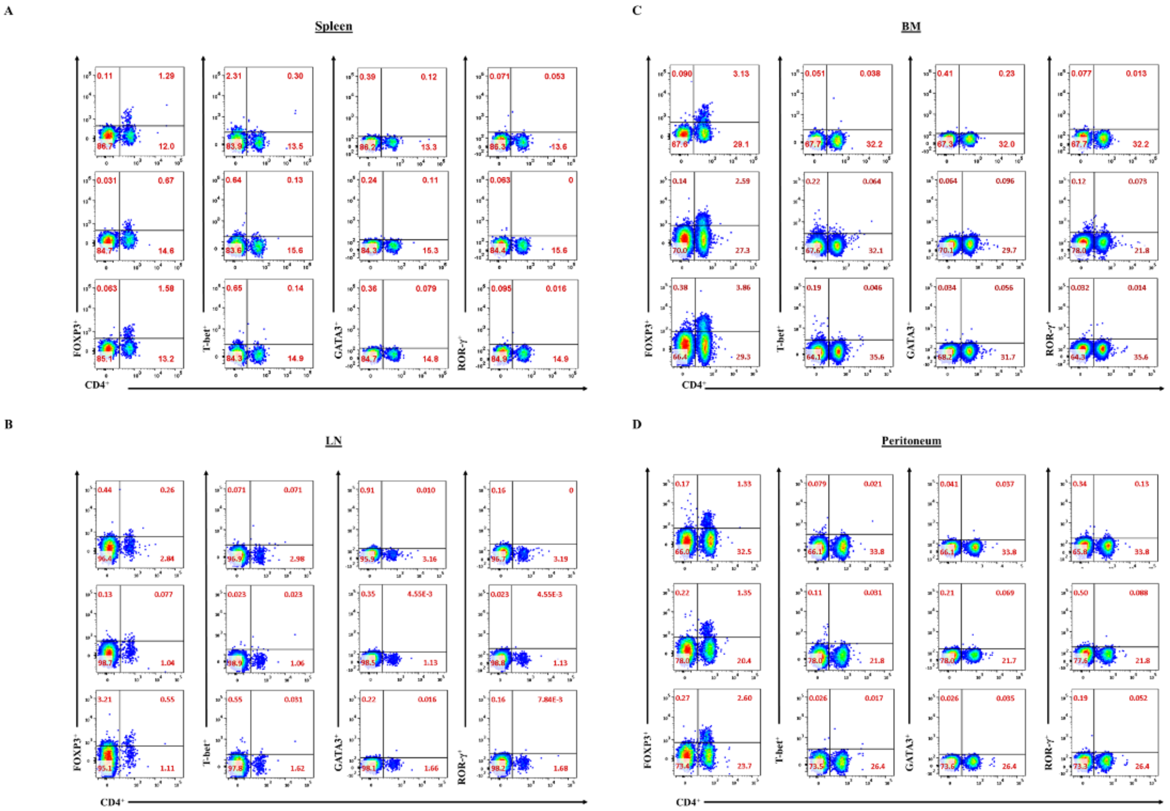

**Supplementary Figure 4**

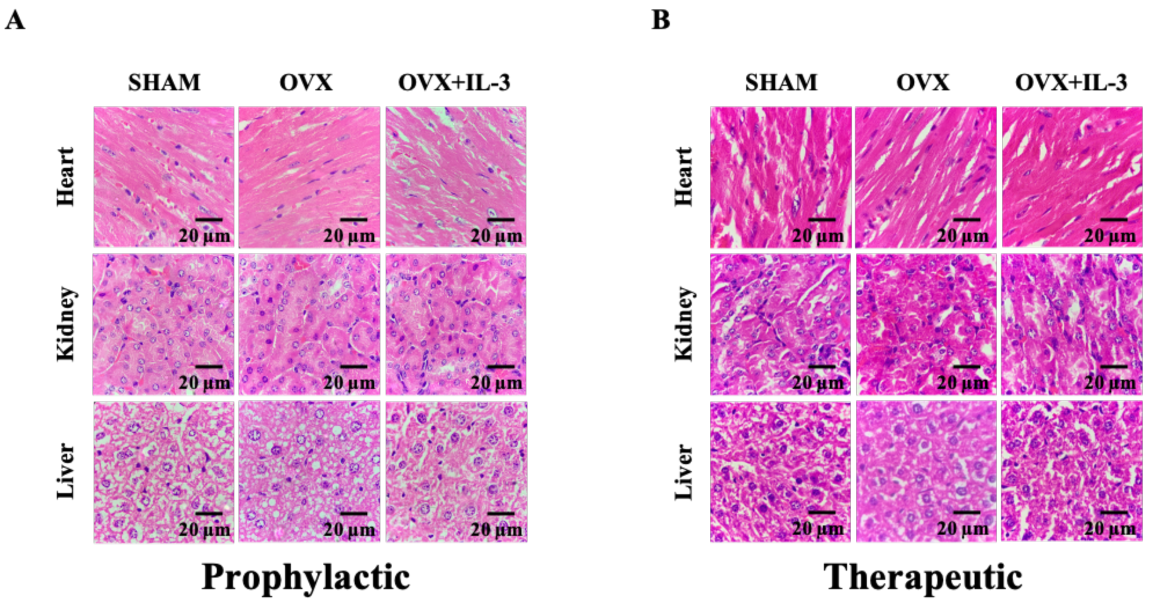

**Table S1: List of primers and sequence for qPCR**

| Gene | Sense primer (5'-3') | Antisense primer (5'-3') |
|---|---|---|
| **Itgb3** | AGAATGCCTGCTTGCCCATGT | TACGGGACACGCTCTGTTTCT |
| **Dc-stamp** | AGAGCTGTTGACTTCCGGG | ATACTTCCAGCCACAAGGGC |
| **Nfatc1** | CGTGGAGAAGCAGAGCACAG | CTTGCACAGGTCTCGGTCAG |
| **Ctr** | TGCCAACCATTATCCACGCCA | TCACAAGCACGCGGACAATGT |
| **Ctsk** | TGCCTTCCAATACGTGCAGCA | TGCATTTAGCTGCCTTTGCCG |
| **Tnfrsf11a** | TTCGACTGGTTCACTGCTCC | CCTCAGAATCCACCGTGCTT |
| **Mmp9** | GTGGTTCAGTTGTGGTGGTG | CCCGCTGTATAGCTACCTCG |
| **Trap** | GGAACTTCCCCAGCCCTTAC | AGGTCTCGAGGCATTTTGGG |
| **Alp** | AAGTTCGCTATGTGCCTTGCC | AGGTGCTTTGGGAATCTGTGC |
| **Ocn** | AGACCTAGCAGACACCATGA | TGCTTGGACATGAAGGCT |
| **Osx** | TGCTTGAGGAAGAAGCTCAC | TTTTGGGGGCTGAAAGGTCA |
| **Runx2** | AGAACTGGGCCCTTTTTCAG | TACGTGTGGTAGTGAGTGGT |
| **Opn** | TCCTTGCTTGGGTTTGCAGTC | TGCCAGAATCAGTCACTTTCA |
| **Col1a** | AAGAATGGCGATCGTGGTGAG | AAGCCACGATGACCCTTTATG |
| **Bmp2** | GAAGTTCCTCCACGGCTTCT | AGATCTGTACCGCAGGCACT |
| **Bmp4** | GAGCCATTCCGTAGTGCCAT | TGGTGTCTTGACAGAAAACAAGG |
| **Bmp6** | AACGCACACATGAATGCCAC | AACCACAAGCTCTCACGACC |

**Table S2. Trabecular bone parameters of DFM under strategy-I (Prophylactic)**

| Parameter | SHAM (n=10) | OVX (n=9) | OVX+IL-3 (n=10) |
|---|---|---|---|
| BMD (g/cm$^3$) | 0.1052 ± 0.003113 | 0.06828 ± 0.003185 | 0.1009 ± 0.003739 |
| BV (mm3) | 0.1779 ± 0.007235 | 0.1056 ± 0.008488 | 0.178 ± 0.01054 |
| BV/TV (%) | 7.413 ± 0.2858 | 4.286 ± 0.2838 | 7.066 ± 0.379 |
| BS (mm$^2$) | 16.76 ± 0.594 | 9.822 ± 0.681 | 14.92 ± 0.7993 |
| BS/TV (mm$^{-1}$) | 6.981 ± 0.2225 | 3.997 ± 0.2371 | 5.926 ± 0.2816 |
| BS/BV (mm$^{-1}$) | 94.47 ± 1.278 | 93.79 ± 2.026 | 84.18 ± 1.215 |
| Tb. Th (mm) | 0.04219 ± 0.0005099 | 0.04208 ± 0.000837 | 0.04648 ± 0.0006119 |
| Tb. N (mm$^{-1}$) | 1.754 ± 0.05827 | 1.016 ± 0.06091 | 1.52 ± 0.07916 |
| Conn. D (mm$^{-3}$) | 198.9 ± 17.97 | 88.62 ± 9.441 | 163.4 ± 13.45 |
| Tb. Sp (mm) | 0.2669 ± 0.00559 | 0.3962 ± 0.02483 | 0.3639 ± 0.01968 |
| Tb. Pf (mm$^{-1}$) | 32.67 ± 0.6326 | 33.02 ± 0.9529 | 26.87 ± 0.6178 |
| SMI | 2.074 ± 0.022 | 2.111 ± 0.02934 | 1.914 ± 0.02258 |

**Table S3. Trabecular bone parameters of PTM under strategy-I (Prophylactic)**

| Parameter | SHAM (n=10) | OVX (n=9) | OVX+IL-3 (n=10) |
|---|---|---|---|
| BMD (g/cm$^3$) | 0.1497 ± 0.00601 | 0.09087 ± 0.005564 | 0.1337 ± 0.005162 |
| BV (mm3) | 0.1932 ± 0.00896 | 0.1184 ± 0.006495 | 0.1891 ± 0.01083 |
| BV/TV (%) | 12.45 ± 0.5653 | 7.397 ± 0.4074 | 11.17 ± 0.5503 |
| BS (mm$^2$) | 15.43 ± 0.6424 | 10.2 ± 0.5128 | 14.45 ± 0.7583 |
| BS/TV (mm$^{-1}$) | 9.943 ± 0.4053 | 6.377 ± 0.3185 | 8.527 ± 0.3599 |
| BS/BV (mm$^{-1}$) | 80.05 ± 0.7081 | 86.41 ± 1.123 | 76.66 ± 1.093 |
| Tb.Th (mm) | 0.04692 ± 0.0003376 | 0.04505 ± 0.0005672 | 0.05011 ± 0.0008449 |
| Tb.N (mm$^{-1}$) | 2.651 ± 0.1137 | 1.646 ± 0.09749 | 2.228 ± 0.1022 |
| Conn.Dn (mm$^{-3}$) | 230.7 ± 13.99 | 140 ± 11.77 | 192.1 ± 13.36 |
| Tb.Sp (mm) | 0.2287 ± 0.006316 | 0.3458 ± 0.02108 | 0.2941 ± 0.01136 |
| Tb.Pf (mm$^{-1}$) | 23.59 ± 0.609 | 29.43 ± 0.9651 | 24.06 ± 0.6318 |
| SMI | 1.766 ± 0.03353 | 2.04 ± 0.05055 | 1.881 ± 0.03183 |

**Table S4: Trabecular bone parameters of DFM under strategy-II (Therapeutic)**

| Parameter | SHAM (n=10) | OVX (n=12) | OVX+IL-3 (n=11) |
|---|---|---|---|
| BMD ($g/cm^3$) | 0.1112 ± 0.002638 | 0.07317 ± 0.004433 | 0.1089 ± 0.003012 |
| BV (mm3) | 0.1486 ± 0.005692 | 0.09041 ± 0.007076 | 0.1612 ± 0.008286 |
| BV/TV (%) | 6.336 ± 0.232 | 4.095 ± 0.3523 | 6.569 ± 0.3109 |
| BS ($mm^2$) | 13.17 ± 0.4118 | 8.414 ± 0.5896 | 13.55 ± 0.6757 |
| BS/TV ($mm^{-1}$) | 5.615 ± 0.165 | 3.81 ± 0.3003 | 5.525 ± 0.2575 |
| BS/BV ($mm^{-1}$) | 88.89 ± 1.065 | 93.9 ± 1.446 | 84.27 ± 1.326 |
| Tb.Th (mm) | 0.04647 ± 0.0005857 | 0.04247 ± 0.0005889 | 0.04711 ± 0.0008541 |
| Tb.N ($mm^{-1}$) | 1.364 ± 0.04957 | 0.9579 ± 0.07575 | 1.401 ± 0.0741 |
| Conn.Dn ($mm^{-3}$) | 126.5 ± 8.607 | 70.13 ± 6.414 | 111.8 ± 7.903 |
| Tb.Sp (mm) | 0.2908 ± 0.004821 | 0.3696 ± 0.02948 | 0.311 ± 0.01055 |
| Tb.Pf ($mm^{-1}$) | 33.12 ± 0.531 | 34.91 ± 0.8601 | 29.84 ± 0.9248 |
| SMI | 2.235 ± 0.02069 | 2.229 ± 0.03142 | 2.121 ± 0.04717 |

**Table S5. Trabecular bone parameters of PTM under strategy-II (Therapeutic)**

| Parameter | SHAM (n=10) | OVX (n=12) | OVX+IL-3 (n=11) |
|---|---|---|---|
| BMD | 0.1573 ± 0.003033 | 0.1017 ± 0.007415 | 0.1483 ± 0.005315 |
| BV | 0.1616 ± 0.005691 | 0.1058 ± 0.009978 | 0.1701 ± 0.008835 |
| BV/TV | 9.978 ± 0.3099 | 6.313 ± 0.6029 | 9.726 ± 0.4178 |
| BS | 11.79 ± 0.351 | 8.582 ± 0.7653 | 12.55 ± 0.6324 |
| BS/BV | 73.13 ± 0.8096 | 81.69 ± 1.084 | 74.03 ± 1.164 |
| BS/TV | 7.287 ± 0.2006 | 5.121 ± 0.4572 | 7.178 ± 0.2822 |
| Tb.Th | 0.05223 ± 0.0006946 | 0.04851 ± 0.0007725 | 0.05201 ± 0.000914 |
| Tb.Sp | 0.3211 ± 0.01494 | 0.4107 ± 0.02934 | 0.3289 ± 0.01417 |
| Tb.N | 1.911 ± 0.05748 | 1.308 ± 0.13142 | 1.87 ± 0.07514 |
| Tb.Pf | 22.91 ± 0.5798 | 28.38 ± 0.9283 | 23.37 ± 0.5079 |
| SMI | 1.879 ± 0.03976 | 2.083 ± 0.05636 | 1.894 ± 0.02345 |
| Conn.Dn | 139.5 ± 7.597 | 114.2 ± 12.58 | 140.3 ± 11.02 |

**Table S6: Cortical bone parameters of femur under strategy-I (Prophylactic)**

| Parameter | SHAM (n=10) | OVX (n=9) | OVX+IL-3 (n=10) |
|---|---|---|---|
| **Cort. BMD** | 1.089 ± 0.005621 | 1.075 ± 0.006717 | 1.086 ± 0.004684 |
| **Cort. BV** | 0.7933 ± 0.0123 | 0.7409 ± 0.02118 | 0.7612 ± 0.01417 |
| **Cort. BV/TV** | 41.59 ± 0.4021 | 38.58 ± 0.6322 | 40.03 ± 0.494 |
| **Cort. Th** | 0.1854 ± 0.002227 | 0.1726 ± 0.003279 | 0.1771 ± 0.002116 |
| **Cort. B. Ar** | 0.75 ± 0.01159 | 0.7006 ± 0.01997 | 0.72 ± 0.01332 |
| **Cort. B. Ar/T. Ar** | 41.72 ± 0.3976 | 38.74 ± 0.617 | 40.16 ± 0.4941 |

**Table S7. Cortical bone parameters of tibia under strategy-I (Prophylactic)**

| Parameter | SHAM (n=10) | OVX (n=10) | OVX+IL-3 (n=10) |
|---|---|---|---|
| **Cort. BMD** | 1.023 ± 0.005614 | 0.9875 ± 0.007286 | 0.9924 ± 0.004477 |
| **Cort. BV** | 0.7873 ± 0.01337 | 0.7476 ± 0.01855 | 0.7937 ± 0.01158 |
| **Cort. BV/TV** | 54.1 ± 0.4296 | 50.27 ± 0.7953 | 51.45 ± 0.5904 |
| **Cort. Th** | 0.1966 ± 0.002334 | 0.1793 ± 0.003624 | 0.1826 ± 0.002197 |
| **Cort. B. Ar** | 0.745 ± 0.01263 | 0.7078 ± 0.01753 | 0.7512 ± 0.01094 |
| **Cort. B. Ar/T. Ar** | 54.26 ± 0.4264 | 50.45 ± 0.7932 | 51.62 ± 0.5897 |

**Table S8. Cortical bone parameters of femur under strategy-II (Therapeutic)**

| Parameter | SHAM (n=10) | OVX (n=10) | OVX+IL-3 (n=10) |
|---|---|---|---|
| **Cort. BMD** | 1.114 ± 0.005688 | 1.074 ± 0.006682 | 1.115 ± 0.003416 |
| **Cort. BV** | 0.9336 ± 0.01475 | 0.7983 ± 0.02076 | 0.8871 ± 0.01088 |
| **Cort. BV/TV** | 48.49 ± 0.4506 | 43.42 ± 0.9105 | 45.38 ± 0.4049 |
| **Cort. Th** | 0.2192 ± 0.001807 | 0.1877 ± 0.004039 | 0.2048 ± 0.001986 |
| **Cort. B. Ar** | 0.8822 ± 0.01391 | 0.7546 ± 0.01957 | 0.8384 ± 0.01027 |
| **Cort. B. Ar/T. Ar** | 48 ± 0.4514.61 | 43.54 ± 0.9096 | 45.49 ± 0.4051 |

**Table S9. Cortical bone parameters of femur under strategy-II (Therapeutic)**

| Parameter | SHAM (n=10) | OVX (n=10) | OVX+IL-3 (n=10) |
|---|---|---|---|
| **Cort. BMD** | 1.052 ± 0.003689 | 1.017 ± 0.007803 | 1.054 ± 0.003337 |
| **Cort. BV** | 0.9268 ± 0.01116 | 0.8217 ± 0.02132 | 0.9395 ± 0.01526 |
| **Cort. BV/TV** | 60.64 ± 0.3886 | 56.21 ± 0.8658 | 59.22 ± 0.6165 |
| **Cort. Th** | 0.2239 ± 0.001729 | 0.1963 ± 0.004385 | 0.2223 ± 0.003561 |
| **Cort. B. Ar** | 0.8768 ± 0.01053 | 0.7777 ± 0.0201 | 0.8888 ± 0.0144 |
| **Cort. B. Ar/T. Ar** | 60.8 ± 0.3884 | 56.38 ± 0.8622 | 59. 37 ± 0.6144 |

**Table S10. Haematological parameters of experimental animals under strategy-I (Prophylactic)**

| Parameters | SHAM (n=8) | OVX (n=8) | OVX+IL-3 (n=8) |
|---|---|---|---|
| RBC Count (X $10^6$/µl) | 5.65 ± 0.15 | 5.52 ± 0.13 | 5.51 ± 0.17 |
| Hematocrit (%) | 33.70 ± 0.89 | 33.42 ± 0.53 | 32.82 ± 0.75 |
| MCV (fl) | 59.78 ± 1.31 | 60.97 ± 2.15 | 59.90 ± 1.63 |
| MCH (pg) | 24.43 ± 0.64 | 23.70 ± 0.79 | 24.34 ± 1.00 |
| MCHC (g/dl) | 40.89 ± 0.85 | 39.04 ± 0.90 | 40.63 ± 1.24 |
| Hemoglobin (Hb) (g/dL) | 13.73 ± 0.23 | 13.02 ± 0.26 | 13.26 ± 0.21 |
| Platelet Count (X $10^3$/µL) | 825.10 ± 25.91 | 806.30 ± 25.23 | 821.90 ± 16.09 |
| Total Leukocyte Count (X $10^3$/µL) | 9.08 ± 0.30 | 9.02 ± 0.29 | 9.10 ± 0.33 |
| Neutrophil (X $10^3$/µL) | 2.04 ± 0.09 | 1.94 ± 0.16 | 1.90 ± 0.13 |
| Lymphocyte (X $10^3$/µL) | 6.69 ± 0.25 | 6.77 ± 0.22 | 6.81 ± 0.23 |
| Monocyte (X $10^3$/µL) | 0.15 ± 0.02 | 0.15 ± 0.01 | 0.18 ± 0.03 |
| Eosinophil (X $10^3$/µL) | 0.19 ± 0.02 | 0.17 ± 0.03 | 0.21 ± 0.03 |
| Basophil (X $10^3$/µL) | 0 | 0 | 0 |
| Neutrophil (%) | 22.50 ± 0.82 | 21.33 ± 1.44 | 20.80 ± 1.07 |
| Lymphocyte (%) | 73.60 ± 0.92 | 75.11 ± 1.40 | 74.90 ± 0.77 |
| Monocyte (%) | 1.70 ± 0.26 | 1.67 ± 0.17 | 2.00 ± 0.30 |
| Eosinophil (%) | 2.10 ± 0.23 | 1.89 ± 0.31 | 2.30 ± 0.30 |
| Basophil (%) | - | - | - |

**Table S11. Hematological parameters of experimental animals under strategy-II (Therapeutic)**

| Parameters | SHAM (n=10) | OVX (n=12) | OVX+IL-3 (n=11) |
|---|---|---|---|
| RBC Count (X $10^6$/µl) | 8.643 ± 0.09 | 8.369 ± 0.12 | 8.916 ± 0.15 |
| Hematocrit (%) | 38.15 ± 0.47 | 36.98 ± 0.70 | 39.66 ± 0.74 |
| MCV (fl) | 44.13 ± 0.13 | 44.2 ± 0.57 | 44.52 ± 0.20 |
| MCH (pg) | 14.89 ± 0.02 | 14.67 ± 0.19 | 14.63 ± 0.072 |
| MCHC (g/dl) | 33.74 ± 0.10 | 33.18 ± 0.13 | 32.86 ± 0.16 |
| Hemoglobin (Hb) (g/dL) | 12.87 ± 0.14 | 12.27 ± 0.23 | 13.04 ± 0.24 |
| Platelet Count (X $10^3$/µL) | 639.9 ± 8.81 | 650.8 ± 21.05 | 762.5 ± 27.95 |
| Total Leukocyte Count (X $10^3$/µL) | 5.49 ± 0.28 | 4.525 ± 0.48 | 6.927 ± 0.85 |
| Neutrophil (X $10^3$/µL) | 2.85 ± 0.16 | 2.125 ± 0.21 | 3.427 ± 0.51 |
| Lymphocyte (X $10^3$/µL) | 2.59 ± 0.20 | 2.417 ± 0.32 | 3.118 ± 0.39 |
| Monocyte (X $10^3$/µL) | 0 | 0 | 0 |
| Eosinophil (X $10^3$/µL) | 0 | 0 | 0 |
| Basophil (X $10^3$/µL) | 0 | 0 | 0 |
| Neutrophil (%) | 39.25 ± 2.19 | 40.42 ± 3.30 | 39.77 ± 1.88 |
| Lymphocyte (%) | 58.8 ± 2.36 | 58.5 ± 3.29 | 57.73 ± 1.60 |
| Monocyte (%) | 0.7 ± 0.21 | 0.5833 ± 0.22 | 0.9091 ± 0.21 |
| Eosinophil (%) | 1.25 ± 0.46 | 0.5 ± 0.15 | 1.591 ± 0.29 |
| Basophil (%) | 0 | 0 | 0 |

**Table S12. Haematological parameters of IL-3 KO animals in comparison with their respective wild type mice**

| Parameters | Reference Values of C57BL6 | Male | | Female | |
|---|---|---|---|---|---|
| | | WT (n=5) | IL-3 KO (n=6) | WT (n=9) | IL-3 KO (n=9) |
| Total Erythrocytes Count ($10^6$/µL) | 7.14-12.20 | 9.18 ± 0.2922 | 8.6 ± 0.1751 | 11.19 ± 0.3238 | 9.856 ± 0.2858 |
| Hemoglobin (g/dL) | 10.8-19.2 | 14.06 ± 0.44 | 12.8 ± 0.2221 | 15.06 ± 0.3823 | 16.72 ± 0.4884 |
| Hematocrit (%) | 37.3-60.0 | 41.18 ± 1.371 | 37.75 ± 0.7632 | 49.49 ± 1.476 | 44.51 ± 1.233 |
| MCV (fL) | 42.7-56.0 | 44.88 ± 0.09695 | 43.88 ± 0.4615 | 44.24 ± 0.09146 | 45.22 ± 0.2314 |
| MCH (pg) | 11.7-16.3 | 15.32 ± 0.08 | 14.88 ± 0.1493 | 14.94 ± 0.08837 | 15.32 ± 0.1176 |
| MCHC(g/dL) | 24.6-34.9 | 34.04 ± 0.1166 | 33.92 ± 0.2242 | 33.8 ± 0.1563 | 33.84 ± 0.1709 |
| Platelet Count ($10^6$/µL) | 285-890 | 622.4 ± 16.66 | 586.3 ±33.67 | 405.1 ± 28.86 | 454 ± 22.35 |
| Total Leucocyte Count ($10^3$/µL) | 4.45-13.96 | 8.68 ± 0.6414 | 8.667 ± 1.054 | 7.1 ± 0.6375 | 5.967 ± 0.6526 |
| Lymphocyte ($10^3$/µL) | 3.24-11.15 | 5.66 ± 0.5144 | 4.917 ± 0.6828 | 4.667 ± 0.4546 | 3.756 ± 0.4289 |
| Monocyte ($10^3$/µL) | 0.15-0.94 | ND | ND | 0.04444 ± 0.03379 | 0.02222 ± 0.02222 |
| Eosinophil ($10^3$/µL) | 0.01-0.42 | 0.28 ± 0.196 | 0.4667 ± 0.3051 | 0.08889 ± 0.04843 | 0.01111 ± 0.01111 |
| Neutrophils ($10^3$/µL) | 0.53-3.09 | 2.74 ± 0.1939 | 3.283 ± 0.4534 | 2.3 ± 0.2466 | 2.156 ± 0.2625 |
| Lymphocyte (%) | 61.26-87.18 | 64.8 ± 2.905 | 56 ± 3.215 | 65.11 ± 1.399 | 62 ± 1.424 |
| Monocyte (%) | 2.18-11.02 | 1.8 ± 0.5831 | 1.667 ± 0.3333 | 2.333 ± 0.527 | 2.556 ± 0.5556 |
| Eosinophil (%) | 0.13-4.42 | 4.6 ± 1.4 | 4.833 ± 2.315 | 2.556 ± 0.5556 | 2.444 ± 0.4444 |
| Neutrophil (%) | 7.36-28.59 | 28.8 ± 1.744 | 37.5 ± 4.507 | 30.11 ± 1.55 | 33 ± 1.225 |
| Basophil (%) | 0.01-1.24 | ND | ND | ND | ND |
| Reticulocytes | 1.8-5.2 | 1.32 ± 0.05831 | 1.217 ± 0.07923 | 2.522 ± 0.1441 | 2.333 ± 0.1764 |

**Table S13. Plasma biochemical indices of IL-3 KO vs. wild type mice**

| Parameters | Reference range | Male | | Female | |
|---|---|---|---|---|---|
| | | WT (n=3) | IL-3 KO (n=3) | WT (n=3) | IL-3 KO (n=3) |
| ALB (g/dL) | 2.5-4.8 | 2.87 ± 0.04 | 2.75 ± 0.06 | 2.96 ± 0.03 | 3.1 |
| ALP (U/L) | 62-209 | 162 ± 9.02 | 104 ± 12.29 | 183.7 ± 9.87 | 106.7 ± 3.71 |
| ALT (U/L) | 28-132 | 67.5 ± 14.55 | 54 ± 14.74 | 64.67 ± 4.37 | 47.67 ± 11.89 |
| MYL (U/L) | 1691-3615 | 1941 ± 14.98 | 1621 ± 119.5 | 1789 ± 81 | 150 ± 69.69 |
| Ca (mg/dL) | 5.9-9.4 | 10.3 ± 0.070 | 10.08 ± 0.12 | 10.3 ± 0.15 | 10.07 ± 0.03 |
| CHOL (mg/dL) | 36-96 | 109.5 ± 2.98 | 79.5 ± 9.042 | 84.67 ± 4.91 | 78 ± 2.08 |
| CREA (mg/dL) | 0.2-0.8 | 0.275 ± 0.02 | 0.2± 0.040 | 0.2 | 0.16 ± 0.03 |
| GLU (mg/dL) | 90-192 | 106.3 ± 8.43 | 109.5 ± 13.07 | 64.67 ± 14.44 | 106.7 ± 7.75 |
| PHOS (mg/dL) | 8.6 | 7.8 ± 0.42 | 7.22 ± 0.370 | 6.1 ± 0.05 | 7.2 ± 0.17 |
| TBIL (mg/dL) | 0.1 | 0.1 | 0.17 ± 0.075 | 0.133 ± 0.03 | 0.1 |
| TP (g/dL) | 3.6-6.6 | 5.9 ± 0.04 | 5.97 ± 0.110 | 6.1 ± 0.05 | 6.13 ± 0.06 |
| BUN (mg/dL) | 18-29 | 30 ± 1.73 | 21.25 ± 1.65 | 25.67 ± 2.66 | 23 ± 2.08 |
| GLOB | NA | 3.02 ± 0.025 | 3.2 ± 0.070 | 3.13 ± 0.08 | 2.96 ± 0.03 |

**Table S14: Trabecular bone parameters of DFM in IL-3 KO mice as compare to OVX mice**

| Parameter | SHAM (n=7) | OVX (n=7) | IL-3 KO (n=7) |
|---|---|---|---|
| **BMD (g/cm$^3$)** | 0.1085 ± 0.003904 | 0.08501 ± 0.005254 | 0.08144 ± 0.005231 |
| **BV/TV (%)** | 6.802 ± 0.3259 | 5.074 ± 0.3524 | 3.504 ± 0.5533 |
| **Tb.Th (mm)** | 0.04205 ± 0.0005052 | 0.04264 ± 0.0007058 | 0.04088 ± 0.002457 |
| **Tb.N (mm$^{-1}$)** | 1.618 ± 0.07948 | 1.192 ± 0.08395 | 0.8417 ± 0.112 |
| **Conn.Dn (mm$^{-3}$)** | 133.9 ± 14.1 | 84.55 ± 9.687 | 54.48 ± 9.297 |
| **Tb.Sp (mm)** | 0.2643 ± 0.004957 | 0.3114 ± 0.01108 | 0.3184 ± 0.01363 |

**Table S15: Trabecular bone parameters of PTM in IL-3 KO mice as compare to OVX mice**

| Parameter | SHAM (n=7) | OVX (n=7) | IL-3 KO (n=7) |
|---|---|---|---|
| **BMD (g/cm$^3$)** | 0.1433 ± 0.00644 | 0.1098 ± 0.006677 | 0.1115 ± 0.006031 |
| **BV/TV (%)** | 10.48 ± 0.4314 | 7.729 ± 0.6718 | 6.376 ± 0.6241 |
| **Tb.Th (mm)** | 0.04606 ± 0.001391 | 0.04658 ± 0.0011 | 0.04975 ± 0.001599 |
| **Tb.N (mm$^{-1}$)** | 2.28 ± 0.08527 | 1.673 ± 0.1635 | 1.298 ± 0.1368 |
| **Conn.Dn (mm$^{-3}$)** | 203.5 ± 15.81 | 148.1 ± 33.91 | 97.62 ± 11.45 |
| **Tb.Sp (mm)** | 0.2402 ± 0.00566 | 0.2974 ± 0.01674 | 0.3432 ± 0.02361 |
